# Towards a best-in-class IgE disruptor for food allergy with Claude-enabled antibody design

**DOI:** 10.64898/2026.11.09.765431

**Authors:** Ezra Erives, Emilio Ferrara, Sean Michael, Nikhil Mallipeddi, Bowen Jing

**Author notes:** These authors contributed equally (Ezra Erives, Emilio Ferrara).

## Abstract

Food allergy affects an estimated 220–250 million people worldwide, for whom the constant risk of a severe reaction weighs heavily on quality of life. Allergic reactions are triggered by the cross-linking of allergen-specific IgE bound to its high-affinity receptor, Fc*ε*RI, on mast cells and basophils, causing degranulation, inflammation and, in severe cases, anaphylaxis. Omalizumab (Xolair), the sole FDA-approved biologic for food allergy, neutralizes circulating IgE; however, it does not effectively remove IgE from Fc*ε*RI, where it can persist for weeks and where even a small residual amount can sustain sensitivity. Here we describe an agentic computational design campaign with Claude toward a disruptive inhibitor antibody that physically removes bound IgE from Fc*ε*RI. Guided by our research team, Claude pursued a biparatopic molecule that pairs a *disruptor arm*, which displaces IgE from Fc*ε*RI, with a non-competing *anchor arm* that holds the molecule on receptor-bound IgE. Of 91 experimentally characterized disruptor designs, 20 showed activity against receptor-bound IgE by competitive bio-layer interferometry, and in a dose-titration assay three variants removed more IgE than omalizumab at therapeutically relevant concentrations (25–400 nM). *De novo* designs for the anchor arm bound IgE at diverse epitopes with apparent affinities of 3.4–235 nM. These subcomponents provide the starting point for further characterization toward a best-in-class IgE disruptor.

## 1 Introduction

Food allergy is an atopic disease characterized by the overproduction of IgE reactive against otherwise harmless environmental or food antigens. Allergen-induced cross-linking of IgE bound to the high-affinity receptor Fc*ε*RI of mast cells and basophils releases preformed and newly synthesized inflammatory mediators, resulting in classical symptoms ranging from moderate urticaria to severe, fatal anaphylaxis (Figure 2b) [1]. In the United States the disease affects up to 8% of children and about 10% of adults, roughly 33 million people in all, with about 40% of affected children reactive to more than one food [2–4]. Until recently, the only option for food allergy patients was strict allergen avoidance and self-injectable epinephrine for accidental exposures [5].

**Figure 1.**
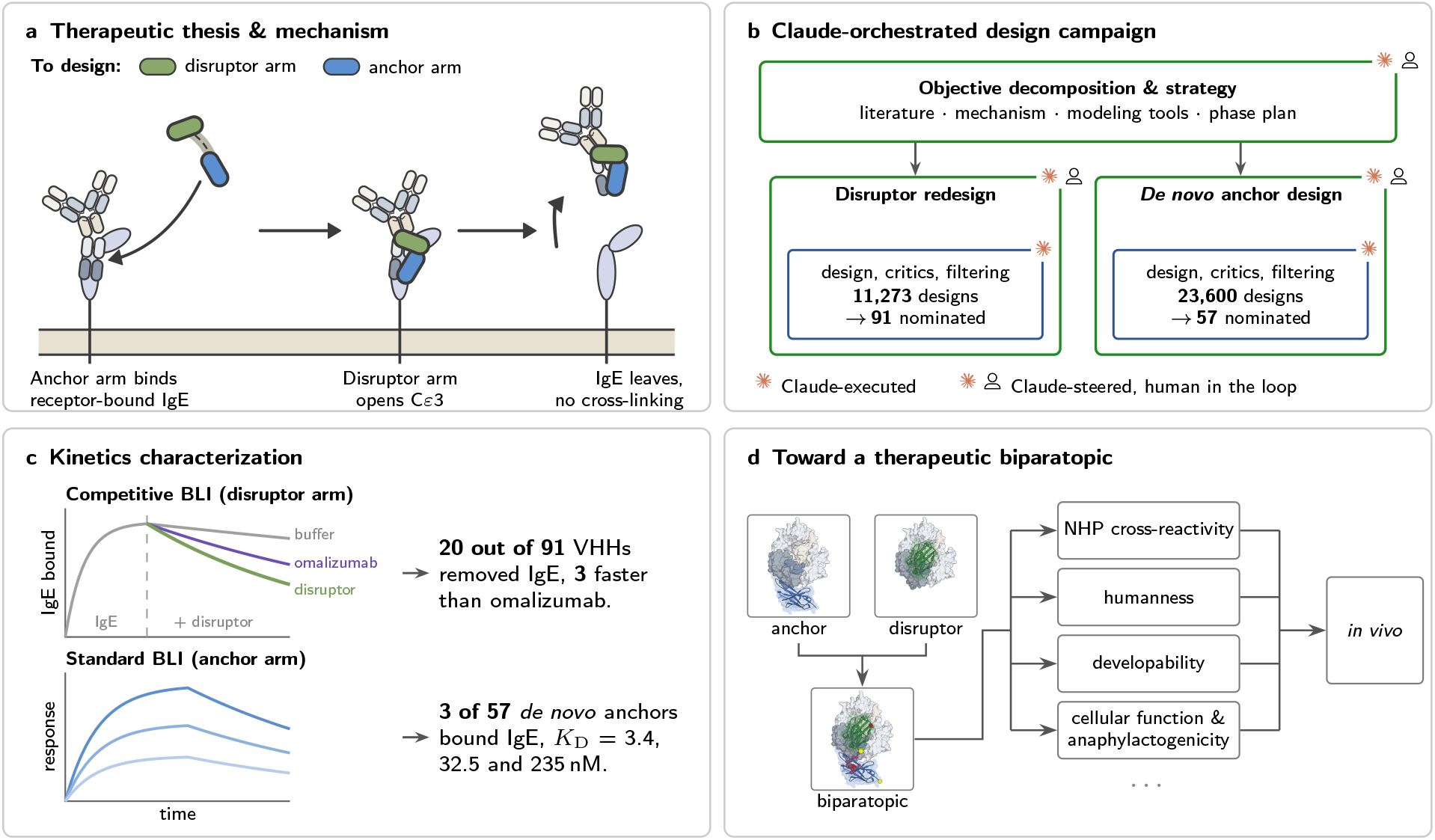
Overview of biparatopic disruptive IgE inhibitor design campaign. **(a)** We sought to design a biparatopic VHH whose anchor arm binds receptor-bound IgE and whose disruptor arm forces IgE into a non-Fc*ε*RI*α*-competent state, so that it departs the receptor without cross-linking (from Figure 2d). **(b)** The design campaign, simplified from Figure 3. Claude decomposed the objective with human input and ran both design phases. **(c)** Characterization by bio-layer interferometry: competitive BLI for the disruptor arm and standard BLI for the anchor arm. **(d)** Judicious pairing of the two arms via geometric analysis will provide the starting point for further *in vitro* and *in vivo* characterization toward a therapeutic biparatopic.

**Figure 2.**
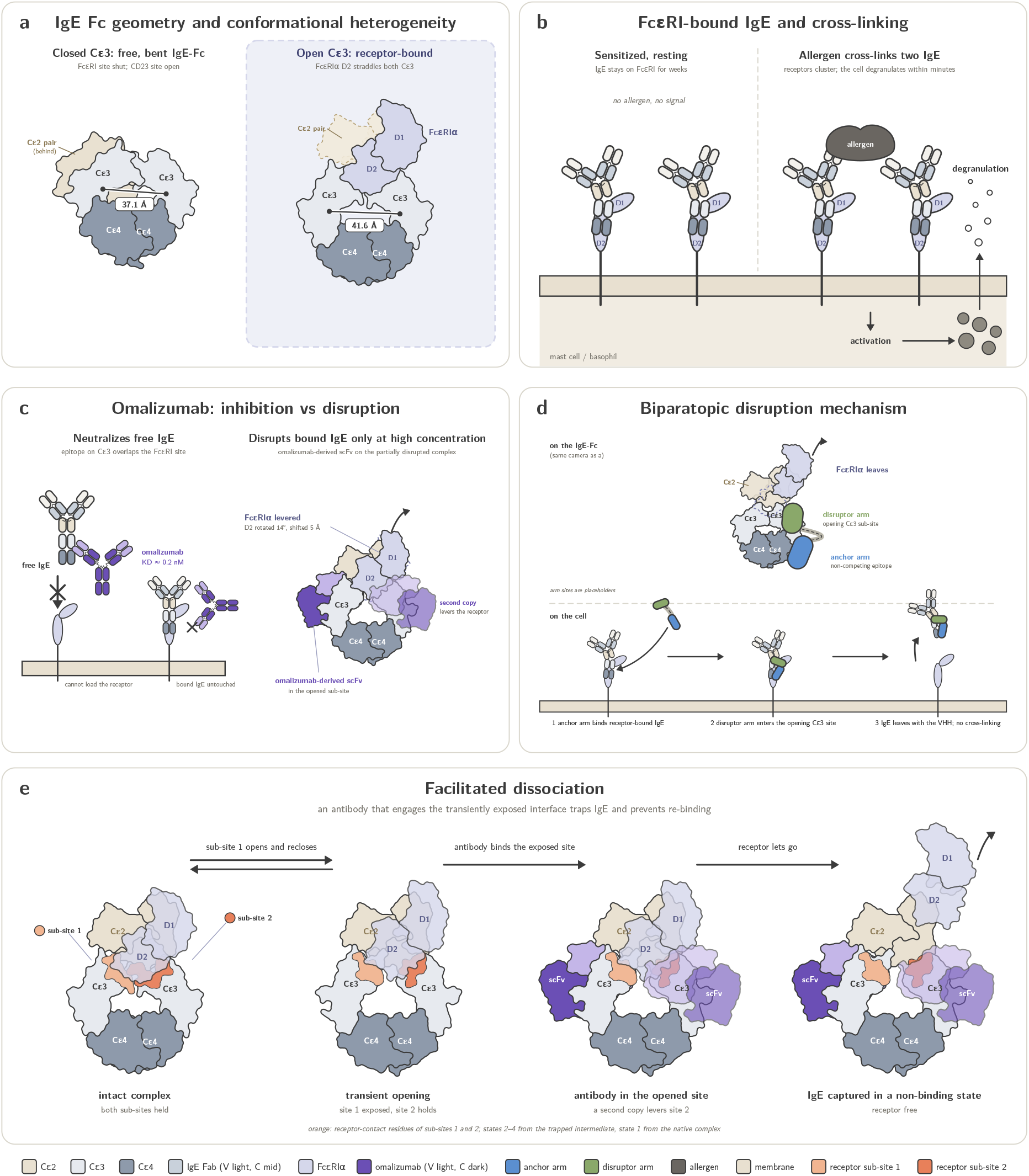
IgE biology and the disruption problem. **(a)** IgE-Fc conformations. Free IgE-Fc (PDB 2WQR) is bent, with the C*ε*2 pair folded back and the C*ε*3 domains closed; receptor-bound IgE-Fc (PDB 1F6A) has the C*ε*3 domains open, straddled by Fc*ε*RI*α* D2. C*ε*3 center-to-center distances are marked; the C*ε*2 pair on the right is from PDB 7SHT (dashed). **(b)** Allergen-induced clustering of IgE:Fc*ε*RI on an effector cell activates the cell and releases granule mediators. **(c)** Omalizumab binds free IgE on C*ε*3 and blocks receptor loading, but its epitope is covered on receptor-bound IgE. Right, a partially disrupted complex (PDB 7SHT). **(d)** Biparatopic disruptor concept. An anchor arm (blue) binds a non-overlapping epitope and holds the molecule on receptor-bound IgE; a disruptor arm (green) enters the C*ε*3 site as it opens and levers the receptor off. Arm positions are illustrative (E3_53 and E2_79 sites, PDB 7MXI). **(e)** Facilitated dissociation of IgE:Fc*ε*RI*α* complex by omalizumab, and antibody variants derived from omalizumab. The IgE:Fc*ε*RI*α* complex opens transiently at one of its two C*ε*3 sub-sites (orange; sub-site 1 light, sub-site 2 dark) while the other holds; an antibody that binds the exposed site blocks re-engagement, the second sub-site releases, and IgE leaves bound to the antibody in a non-binding state. State 1, native complex (PDB 1F6A, C*ε*2 pair from 7SHT); states 2–4, the trapped disruption intermediate (PDB 7SHT) with an omalizumab-derived scFv in the opened sub-site 1 and a second copy levering Fc*ε*RI*α* at sub-site 2 (14^*°*^, 5 Å), shown without the antibody (2) and without the receptor (4). Structures superposed on the C*ε*4 pair, one camera; domains drawn as heavy-atom outlines.

The fundamental pathway of food allergy is the interaction of allergen-specific IgE with the high-affinity receptor Fc*ε*RI of mast cells and basophils, one of the strongest protein-receptor interactions in human immunology. IgE binds Fc*ε*RI*α* with an affinity near 10^*−*10^ M and dissociates so slowly that mast cells and basophils stay fully sensitized whether or not allergen is present [6–8], supported by continuous IgE production by B cells [9–11]. While free IgE has a serum half-life of about two days, receptor-bound IgE persists in human skin for weeks to months [9, 12–14]. Additionally, mast cells and basophils are exquisitely sensitive to IgE cross-linking. For reference, a basophil carries some 10^5^ Fc*ε*RI, yet only a few thousand allergen-specific IgE molecules, one or two per cent of the receptor pool, give a half-maximal response, and only a few hundred are enough to initiate secretion [15–17]. Therefore, food allergen reactivity is the result of a baseline state of fully sensitized and highly reactive mast cells and basophils driven by continuous allergen-specific IgE production by B cells.

Over the past decade, the management and treatment of food allergy have been revolutionized. Most notably, in 2024 omalizumab (Xolair), a humanized anti-IgE antibody, became the first FDA-approved medication to reduce allergic reactions to multiple foods [5, 18, 19]. In the OUtMATCH trial, 16 to 20 weeks of omalizumab allowed 67% of patients with multi-food allergy to tolerate 600 mg of peanut protein, against 7% on placebo [18]. This is a major advance for the treatment of food allergy, yet it is far from completely addressing the disease. Only 42% of treated patients completed the full 6,044 mg cumulative challenge for two of their foods and 24% for all three. Furthermore, 14% could not consume even 30 mg of peanut after the full 16 to 20 weeks of treatment [18]. The treatment regimen is also burdensome. Because omalizumab neutralizes IgE stoichiometrically, the dose is set by body weight and total IgE. Patients must take 75 to 600 mg of drug by injection every two or four weeks, with no established dosing for patients whose body weight and/or total IgE lie outside the approved label, excluding a significant population. Furthermore, because omalizumab does not alter IgE production, free IgE returns after discontinuation of drug, requiring patients to be treated indefinitely [5, 19]. Omalizumab therefore protects against accidental exposure, but patients are advised to continue avoiding their allergens and must carry epinephrine [5, 19].

Omalizumab nonetheless represents a breakthrough in the treatment of food allergy and validates IgE as the central mediator of the disease. Its limitations derive from the biology of the IgE:Fc*ε*RI*α* interaction and from the development goals set for omalizumab in the 1990s (Figure 2) [20, 21].

IgE exists in two forms, free in the circulation and bound to Fc*ε*RI on the cell surface, and at therapeutically relevant doses omalizumab neutralizes only the free form (Figure 2c). Because IgE triggers reactions through clustering of the bound form, neutralizing free IgE disarms the cell only as fast as its existing complexes turn over: basophils lose most of their surface IgE within about ten days, but Fc*ε*RI-positive mast cells in skin are unchanged after one week and reduced only after ten [22–24]. The medicine with the greatest potential for patients with food allergy would therefore not only neutralize free IgE but also remove the IgE already bound to effector cells (Figure 2).

## 2 Neutralizing and disruptive IgE inhibitors

Omalizumab neutralizes free IgE by binding to the IgE C*ε*3 chain at two symmetrically related epitopes, and in a manner which both directly overlaps with the bound conformation of Fc*ε*RI*α*, and which additionally prevents the folded-back conformation of the C*ε*2-domain believed to stabilize the IgE:Fc*ε*RI*α* complex (Figure 2a) [20, 21, 25, 26]. For these same reasons, it was originally thought omalizumab was unable to bind IgE once it is already bound to Fc*ε*RI*α*. However, subsequent studies of omalizumab after its initial development discovered that it could in fact disrupt IgE:Fc*ε*RI*α* complexes, but only at concentrations far above those reached in patients. In this case, transient partial dissociation on the part of Fc*ε*RI*α* allows omalizumab to occupy its partially overlapping epitope on C*ε*3, thereby preventing rebinding to Fc*ε*RI*α* (Figure 2e) [25]. Indeed, at micromolar concentrations omalizumab has been shown to accelerate the dissociation of pre-formed complexes and to remove IgE from basophils over hours. Omalizumab’s 50% disruption capacity (DC50) is *∼*200 *µ*M (SPR) for bound IgE disruption compared to KD *≈*.2 nM for free IgE inhibition [25, 27].

### Orthosteric disruptive inhibitors

In parallel to this work, designed ankyrin repeat protein (DARPin) E2_79 was discovered and found to inhibit mediator release more effectively than omalizumab in cellular assays [28, 29]. In 2012, a fluorescence assay developed to classify IgE inhibitors demonstrated that E2_79 disrupted not as a classical competitor of the Fc*ε*RI*α* epitope, but rather through a modeled steric overlap of only 400 Å^3^ in comparison to omalizumab’s modeled value of 1200 Å^3^, suggesting that large overlap may not be necessary and may even be counter-productive to facilitating disruption [8, 25, 30]. Subsequently, E2_79 was shown to remove surface IgE from allergic-donor basophils within minutes, to abolish IgE-dependent activation, and to protect mice carrying human Fc*ε*RI*α* from passive cutaneous anaphylaxis [27]. The DARPins that revealed facilitated dissociation, however, made a difficult starting point for a medicine, limited by immunogenicity, developability, and no approved precedent for the scaffold [27, 31]. Based on this work, the academic groups behind the DARPins turned to engineering omalizumab itself to establish disruptive potency [32, 33]. They evolved variants that strip humanized mast cells at a few hundred nanomolar, and a cryo-EM structure of a variant bound to a partially dissociated complex confirmed steric facilitated dissociation as the mechanism of disruption [32]. Elbow-flexible descendants of these variants desensitize mice within 36 hours of a single dose, where omalizumab matches vehicle [33]. Excellergy, a biotechnology company founded on this work, has now entered phase 1 clinical trials [33, 34].

### Allosteric disruptive inhibitors

Concurrently, the biotechnology company Ablynx isolated single-domain antibodies (VHHs) against IgE from an unbiased llama immunization campaign, and found several that not only blocked IgE binding to Fc*ε*RI but also displaced receptor-bound IgE from cells [35–37]. The mechanism of one of these VHHs, sdAb026, was resolved later: it binds the CD23 (Fc*ε*RII) site at the C*ε*3–C*ε*4 junction and traps the closed conformation that cannot bind Fc*ε*RI*α*, representing a second route to disruption that acts from a distal epitope rather than at the interface [38]. In cellular assays, sdAb026 strips IgE from basophils within minutes.

### Biparatopics

Together, the orthosteric and allosteric disruptive lineages have established two distinct mechanisms for driving disruption of IgE:Fc*ε*RI*α* complexes: steric facilitated dissociation, in which a binder inserts at a transiently exposed sub-site of the receptor interface and competes the receptor off, and allosteric conformational trapping, in which a binder at the distal CD23 (Fc*ε*RII) site locks C*ε*3 in the closed conformation that cannot hold the receptor [8, 38]. In both cases, and regardless of modality, such monovalent disruptors require concentrations of high hundreds of nanomolar to micromolar on cells to mediate disruption. To significantly improve disruptive potency, biparatopic constructs have been generated by combining a monovalent *disruptor arm* with a second *anchor arm* against a non-competing epitope. Fusing E2_79 to a DARPin that recognizes receptor-bound IgE at a distal C*ε*3–C*ε*4 surface improved efficiency about 100-fold [27, 39]. Furthermore, the knob-in-hole IgG1 Fc-fusion, KIH_E07_79, brought the half-maximal disruptive concentration on whole-blood basophils to 2 nM, roughly 500-fold the potency of E2_79, and cleared IgE from basophils and tissue mast cells within nine hours *in vivo* [30, 39, 40]. Biparatopic VHHs have similarly demonstrated significant disruptive potency gains by pairing sdAb026 with non-competing anchoring VHHs [41]. Biparatopic constructs exhibit improved disruption because of an avidity effect for the IgE:Fc*ε*RI*α* complex. The non-competing arm binds receptor-bound IgE and anchors the molecule to the IgE:Fc*ε*RI*α* complex. This, in turn, holds the disruptive arm at a high local concentration, ready to act whenever the complex transiently adopts the state it recognizes (Figure 2d) [27, 39, 41].

### Anaphylactogenicity

The same biparatopic mechanism whose avid *cis*-binding of a single receptor-bound IgE molecule enables significantly higher disruptive potency also carries the risk of *trans*-binding of two distinct receptor-bound IgE molecules, which can lead to cross-linking and anaphylaxis [32]. As a clinical example, the development of omalizumab variant HAE1 was discontinued after adverse events in two participants during a phase 2 trial. Although the nature of the events has not yet been disclosed, cellular assays demonstrated HAE1 could activate basophils from some donors, attributed to longer dwell time on intact IgE:Fc*ε*RI*α* complexes [32]. The flexible binding geometry of biparatopics demands that significant consideration be taken so as to avoid this unwanted cross-linking. While initial experimental validation suggested that KIH_E07_79, for example, was non-activating [39, 40], subsequent testing in additional donors reported a high degree of anaphylactogenic activity [41]. Further characterization has strongly implicated the arm geometry, linking moiety (tandem or Fc-fusion) and orientation, as well as the affinity and kinetics of the arms themselves, in determining the anaphylactogenicity [39–41]. Hence, a truly best-in-class disruptive IgE inhibitor must engage receptor-bound IgE strongly enough to disrupt the complex, yet be physically unable to bridge two IgE molecules on the same cell.

## 3 A computational design campaign toward a best-in-class IgE inhibitor

Potent and safe disruption of bound IgE calls for precise control over molecular geometry. Conventional antibody discovery via display or immunization provides little control of the binding epitope or pose, making complex therapeutic mechanisms such as IgE disruption challenging to target. On the other hand, biomolecular generative models can specify or predict the geometry before any molecule is made, enabling rational optimization or selection. We hypothesized that a computational design campaign leveraging modern biomolecular generative models, directed by Claude [42], and informed by prior literature and rational epitope and pose selection could accelerate the path towards a best-in-class IgE inhibitor: one able to inhibit free IgE with high affinity, disrupt IgE:Fc*ε*RI on allergic effector cells with potency at therapeutically relevant doses, and do so without cross-linking and activating effector cells.

We set out to design a biparatopic antibody with two functional VHH arms, one that disrupts the IgE:Fc*ε*RI*α* interaction and one that anchors the molecule to receptor-bound IgE at a geometrically compatible epitope. Upon a Claude-assisted review of existing literature, we opted to pursue (1) localized redesign of sdAb026, an existing VHH with observed disruptive activity and known IgE-bound structure [38, 43]; to obtain our disruptor arm; and (2) *de novo* design of an anchor arm to diversify and control the anchor arm geometry. We ultimately identified three disruptor variants that are comparable in activity to sdAb026 and which are more potent than omalizumab at therapeutically relevant concentrations. We also designed and characterized *de novo* anchors with single-digit nanomolar affinity for IgE. These results represent the first of many critical steps in developing a complex antibody therapeutic like a biparatopic disruptive IgE inhibitor, and provide the critical modular subcomponents for continued therapeutic optimization.

### 3.1 Claude-steered decomposition and orchestration of the design task

The design campaign was largely driven by interaction with Claude. First, in conjunction with our research team, Claude orchestrated high-level design planning and decomposed the stated objective into distinct work phases. Second, it proceeded with near-autonomous execution of these work phases, supported by our research team throughout, as Claude has recently done for *de novo* binder design against diverse targets [42]. In total, the campaign consisted of a total of 22 distinct phases of autonomous execution, of which 15 were primarily devoted toward computational design. We provide a high-level summary of the workflow in Figure 3.

**Figure 3.**
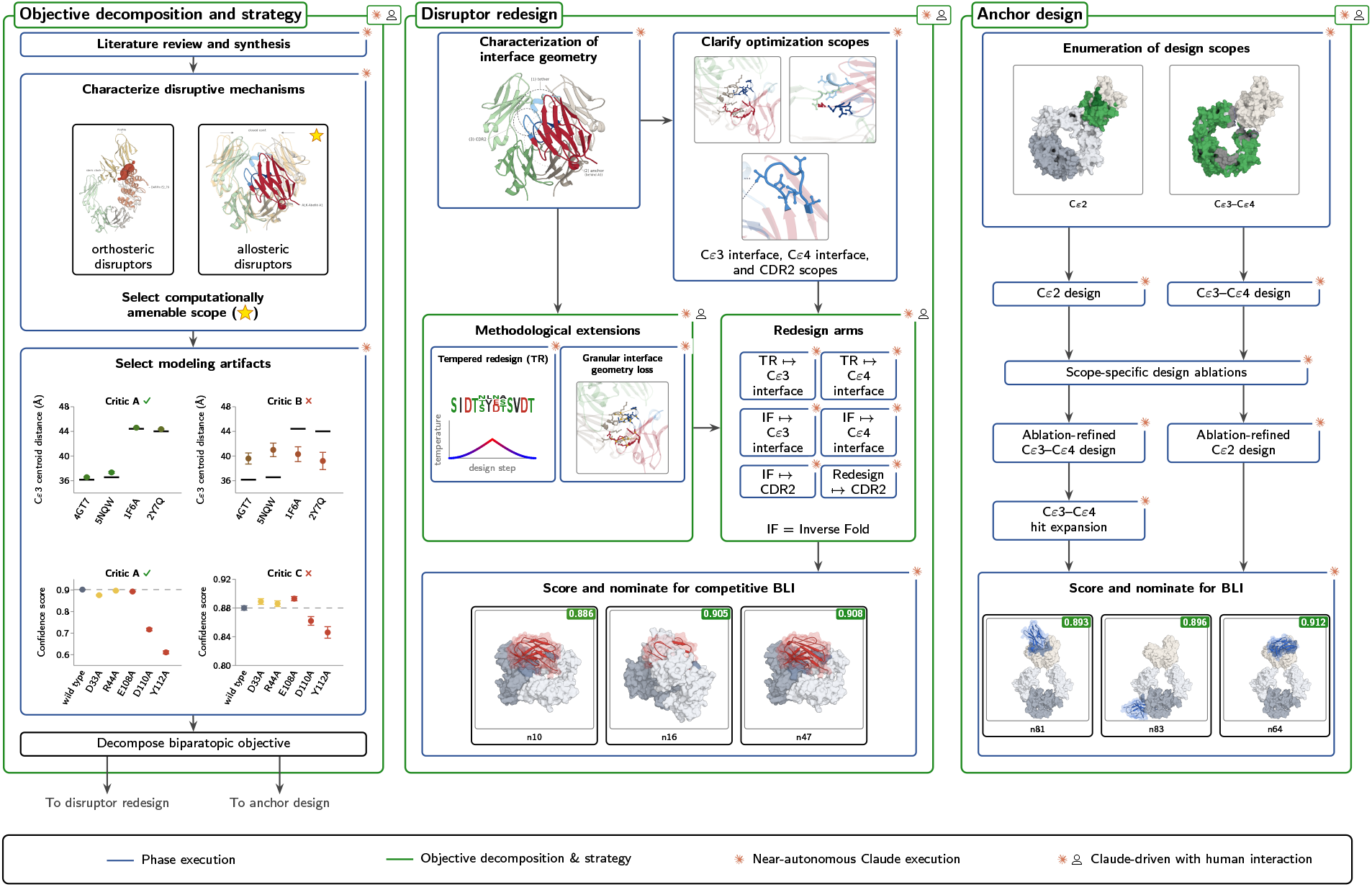
Claude-steered decomposition of the design task. Objective decomposition and strategy, disruptor redesign and de novo anchor design as three phase columns; blue boxes are near-autonomous Claude phase execution, green frames Claude-driven objective decomposition with human interaction.

To initiate the design campaign, Claude performed an extensive review of characterized disruptive approaches from both academic and preclinical work, and grouped these into two salient mechanistic classes: First, **orthosteric disruptors**, including the monovalent and biparatopic lineages extending DARPin E2_79 [8, 27], affinity-evolved, disruption-selected omalizumab variants [32, 33, 39], and even omalizumab it-self at sufficiently high concentration [25, 26], act by exploiting small, transiently freed steric overlap with Fc*ε*RI*α* to gain occupancy and subsequently drive dissociation. Second, **allosteric disruptors** act as CD23 function homologs [44], displaying selective affinity for the non-receptor-competent closed conformation of IgE Fc C*ε*3–C*ε*4, and which includes examples such as sdAb026 [38]. Claude then proceeded to analyze the amenability of each class to computational redesign and eventual inclusion within a biparatopic construct, identifying key functional and structural considerations for each class critical to facilitating disruption and whose accurate recapitulation would be required on the part of the computational model employed.

Toward this end, Claude identified experimentally derived, publicly available data to be used as a means of directly and quantitatively gauging function-specific modeling performance. For example, Claude performed structure prediction on a number of campaign-relevant resolved structures, including both the orthosterically disrupted intermediate state induced by the clone 7 omalizumab variant (PDB 7SHT) [32, 45], and the disruptive binding mode of sdAb026 (PDB 5NQW) [38, 43], observing uniformly high accuracy over both classes in identifying the open/closedness of C*ε*3–C*ε*4, but displaying higher confidence for the allosteric disruptors. Such observations, the existence of an experimentally validated disruptor in VHH sdAb026, and the existence of a solved structure by which to guide redesign, resulted in a final judgment to emulate the allosteric rather than orthosteric disruption mechanism.^1^Finally, to better inform the choice of co-folding models for use in downstream optimization, Claude identified experimentally validated loss-of-function mutants of sdAb026 [38], and used these as the basis of a calibration mechanism by which to select two distinct co-folding models whose confidence most accurately recreated the disruptive rank-ordering of wild type against the disclosed mutants. Claude further clarified the functional specification of the desired biparatopic into respective designs for both arms and a necessary linking moiety. Together with key structural or functional insights, it decomposed the design campaign into subsequent phases. The overall flow of this campaign-planning stage is summarized in Figure 3, left.

### 3.2 Disruptor redesign

To initiate disruptor redesign, Claude began by analyzing the interface geometry of the resolved sdAb026 binding mode to IgE C*ε*3–C*ε*4 in PDB 5NQW [38, 43], identifying three scopes within which to conduct targeted, structurally localized redesign. These scopes include a C*ε*4, receptor-distal interface, mediated by both part of the CDR3 loop and a collection of framework regions (**C***ε***4 interface scope**), a C*ε*3, receptor-proximal interface mediated by CDR1 and CDR3 (**C***ε***3 interface scope**), and a third scope consisting of the interface-distal CDR2 loop (**CDR2 scope**). A fourth scope consisting of CDR1 and CDR3 residues which do not participate in the interface as contacts but which can be inferred to play supporting, interface-stabilizing roles, was advocated for by Claude and carried through the initial inverse-folding and tempered-redesign rounds; however, we opted to drop it from further campaign phases so as to diversify resources away from the binding interface toward the distal CDR2 scope instead. More details on the choice of scopes can be found in Appendix A.1, and a brief visual overview is included in Figure 3.

Claude then elected to split subsequent disruptor redesign efforts into six distinct sub-phases, each consisting of a choice of redesign scope and redesign paradigm. For both the C*ε*3 interface and C*ε*4 interface scopes, Claude opted to perform targeted mutagenesis so as to obtain double, triple, and quadruple mutants with superior disruptive effect. To do so, two primary methods were employed: inverse folding of randomly sampled sub-scopes [46], as well as **tempered redesign**, a purpose-built, gradient-based mutagenesis approach not requiring prior specification of mutation scope. Claude enabled real-time conceptualization, implementation, and subsequent validation and deployment of this key methodological extension, and more broadly meticulously tailored design workflows (both open and internal) to campaign-specific contexts. Finally, both inverse folding and gradient-based redesign were utilized against the CDR2 scope.

At the completion of the round, Claude refolded all designs using the scoring models nominated in Section 3.1, ranked by 10-seed mean binder-pair ipTM, and nominated thirteen distinct designs from each of the four C*ε*3 interface and C*ε*4 interface design scopes, accompanied by an additional forty designs from the CDR2 scope, for subsequent experimental validation. A more detailed overview of the round phasing can be found in Figure 3 (middle column), and a summary of results is reported in Figure 4. We refer the reader to Section 3.2 for a more extensive treatment of each phase mentioned above, and to Appendix C for further details on tempered redesign.

**Figure 4.**
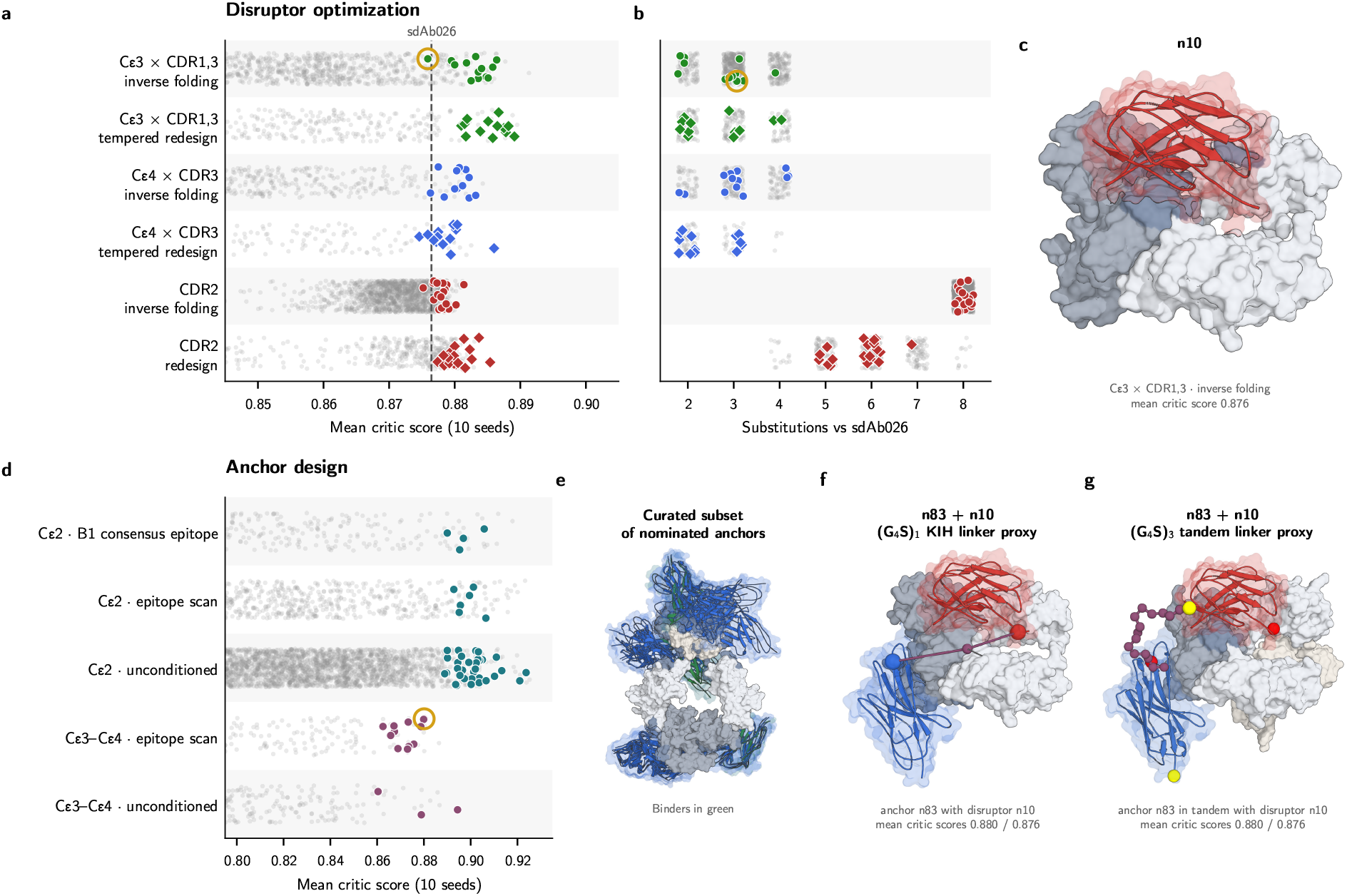
Design campaign results. (a) disruptor redesign (a-c): every scored design per arm (gray) and the nominated designs (colored) by (a) mean critic score over ten seeds and by (b) substitution count against reference sdAb026 (b); dashed line, sdAb026; (c) right, nominee n10 on the C*ε*3–C*ε*4 dimer. Anchor design (d-g): (d) the same strip per design round; (e) a curated subset of nominated anchor variants on both C*ε*2 and C*ε*3–C*ε*4; (f) a (G4S)2 KIH linker proxy applied to disruptor variant n10 and anchor n83, and (g) a (G3S)3 tandem linker proxy applied to the same combination.

### 3.3 De novo anchor design

The anchor arm was designed *de novo* against two surfaces that remain exposed on receptor-bound IgE (Figure 2d). The first is the C*ε*2 domain pair, the epitope class of the VHH anchors that improved sdAb026 in biparatopic format [41]. The second is the C*ε*3–C*ε*4 surface recognized by E3_53, the anchor of the first biparatopic DARPin [27, 39]. Claude first refolded the published C*ε*2-binding VHHs [41] with the selected co-folding critics to check whether the models placed them on a consistent epitope, and used the consensus footprint as the first target.

Anchors were designed by backpropagation through the structure predictor from a random start, over all three CDRs, on twelve clinical VHH frameworks (Table 2). Three phases of design were run on C*ε*2: one against the consensus epitope, one against randomly drawn epitope patches across the domain, and one without epitope conditioning (Appendix B.2). On C*ε*3–C*ε*4, two phases were run: the first with randomly drawn epitope patches as for C*ε*2 and with an additional avoidance loss penalizing contact with the Fc*ε*RI*α* and sdAb026 footprints, as well as a subsequent, completely unconditioned run without epitope conditioning. Both rounds were subsequently followed up by an iterative refinement scheme based on inverse folding [46–49]. In total, 23,600 unique designs were scored in 1,310 H100-hours, and 57 were nominated, 42 on C*ε*2 and 15 on C*ε*3–C*ε*4 (Figure 4, Table 3). We refer the reader to Appendix B for a more technical discussion, to Figure 3 (right column) for a broad overview of the round, and to Figure 4 for results for the round regarding plate nomination for subsequent experimental validation.

**Table 1.**
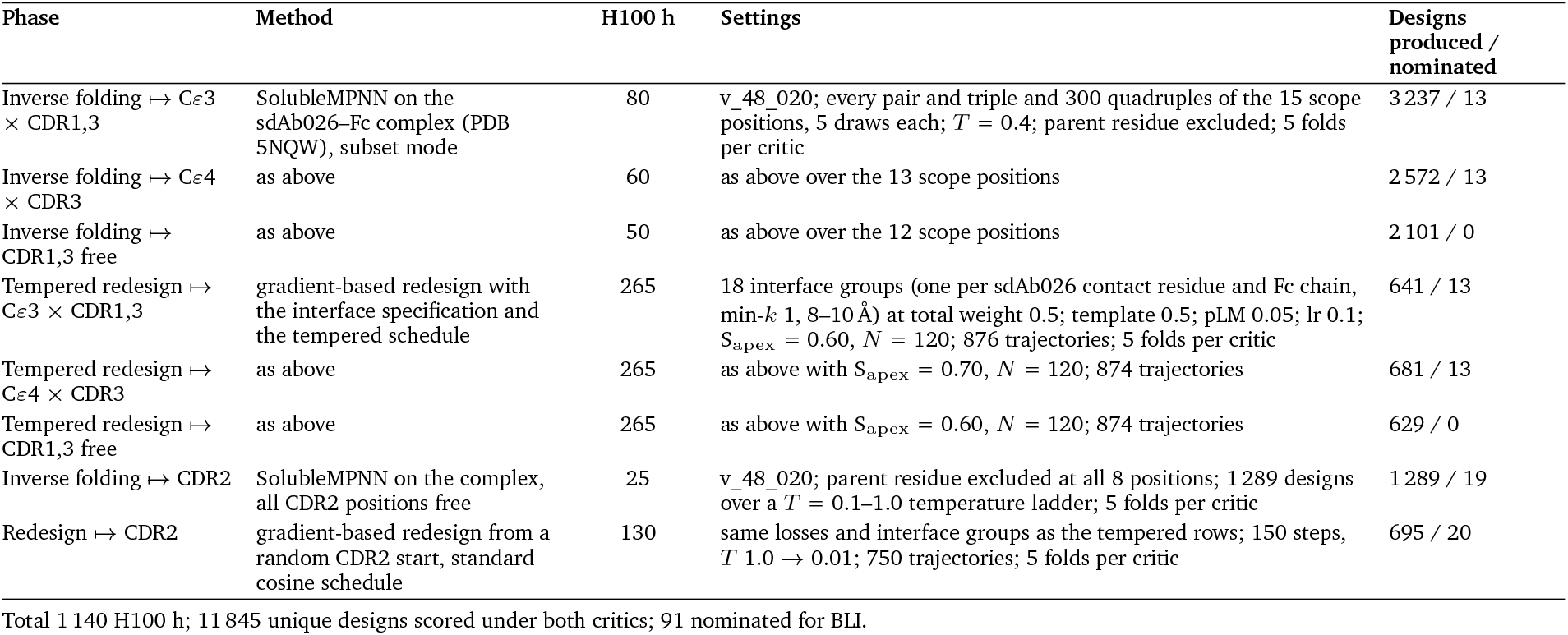
Disruptor optimization: rounds of computation.

**Table 2.**
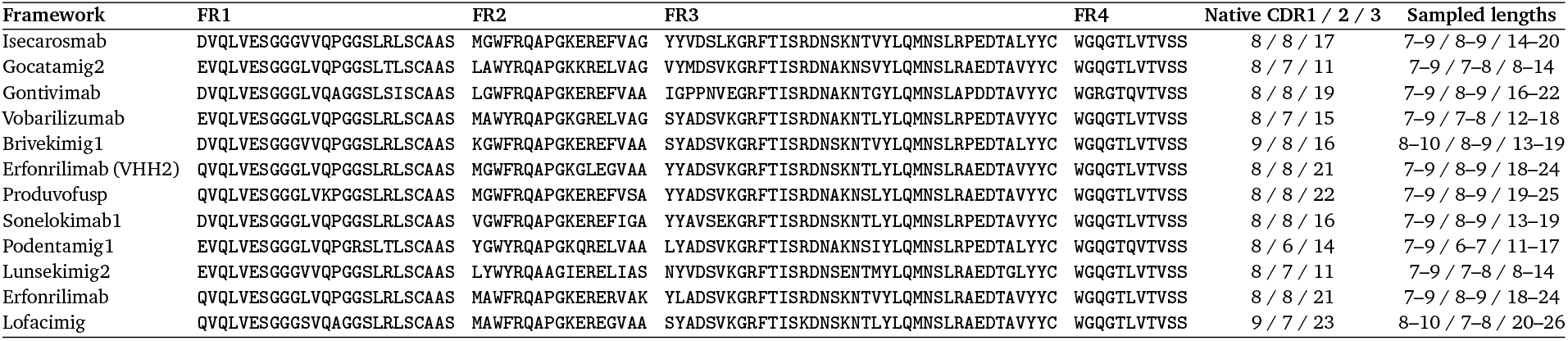
VHH frameworks used for de novo anchor design. Framework regions FR1–FR4 (IMGT) of the twelve therapeutic VHH scaffolds; the three CDR loops between them are designed from random logits with lengths drawn uniformly from the sampled windows (CDR1 / CDR2 / CDR3). Native loop lengths are reported for reference.

**Table 3.**
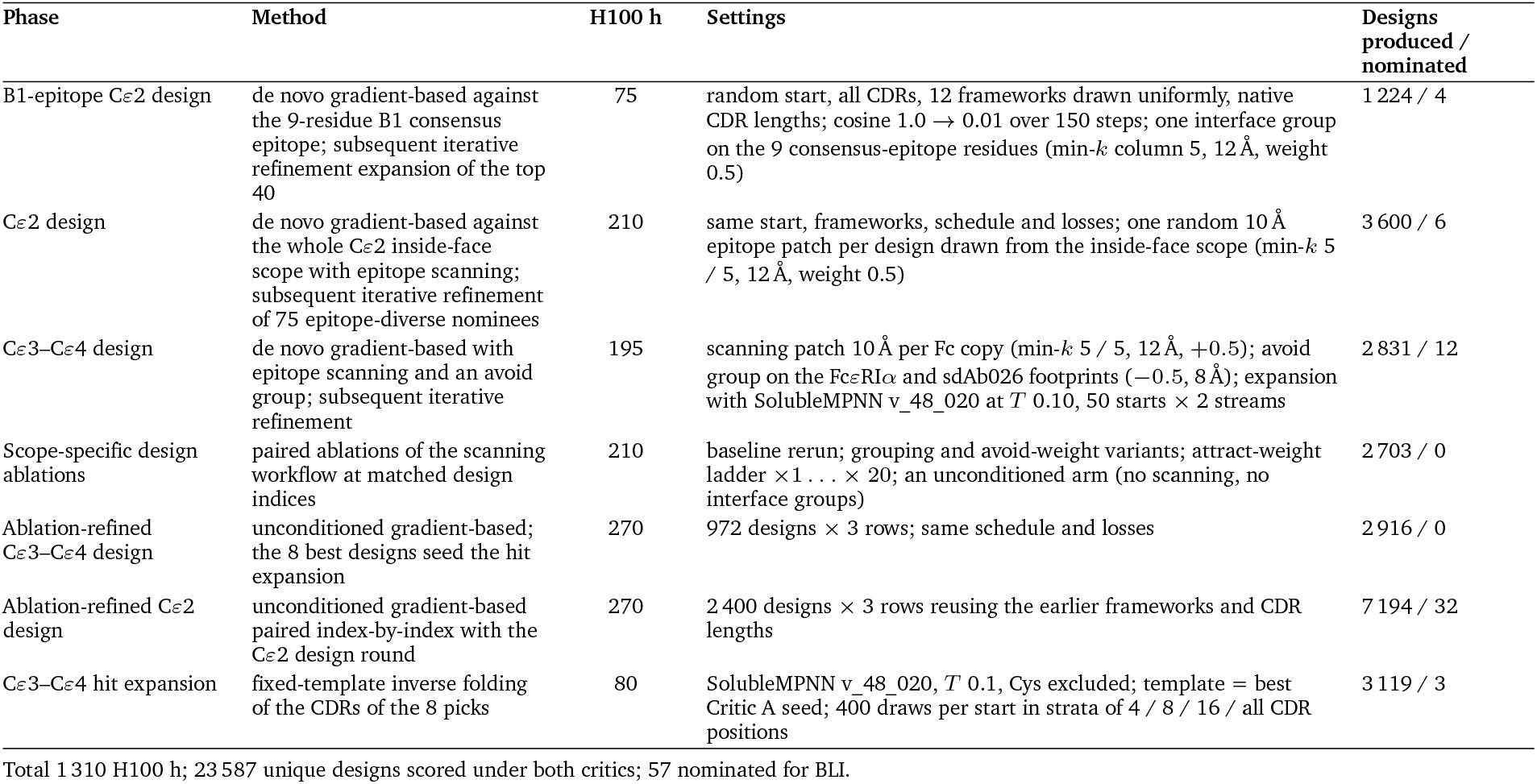
Anchor design: rounds of computation.

### 3.4 Biparatopic assembly

In the final phase of the design campaign, we asked Claude to use the computational results characterizing the disruptive potency and binding affinity of the designs from Section 3.2 and Section 3.3, together with model predictions of pose and binding geometry to assemble promising biparatopics. As discussed in Section 2, the choice of linking moiety varies among precedented disruptors, and within a larger design space consisting of both single-chain tandem linkers, such as the (G_4_S)_2_ linker found in biparatopic DARPin bi53_79 [27], and bulkier, knob-in-hole (KIH), IgG fusion constructs [39, 41]. To further narrow the design mandate, we requested that Claude limit its search to encompass only tandem (G_4_S)_*k*_ linkers for *k∈ {*3, 4, 5, 6, 7, 8 *}*. Despite the limited search space, the dimer symmetry of IgE Fc, combined with the labile nature of C*ε*2, ensures that geometric characterization of the likely cis-binding mode of a putative biparatopic remains a difficult task.

A long and flexible linker is a serious liability for an IgE-targeting biparatopic. The desired biparatopic effect by which an anchor arm binding raises the effective local disruptor arm concentration so as to increase the disruptive potency of the molecule relies on a shorter, rather than longer, linker. The role of linker length, and of linker and arm geometry more generally, is widely implicated as playing a dominant role beyond affinity and kinetics in the anaphylactogenic liability of a particular construct [39, 41]. Indeed, a longer linker and more flexible geometry can accommodate trans binding of distinct receptor-bound molecules, in turn leading to stable bridging, activation, and eventually, anaphylaxis.

With this context present, Claude proceeded to split the task in two: First, Claude conducted a systematic enumeration of potential IgE Fc C*ε*2–C*ε*4 conformations with bound Fc*ε*RI*α* by identifying all deposited structures in the PDB with these chains present (2Y7Q, 7SHT, 8YWA). For each putative pair of anchor and disruptor, all structures were enumerated along with all combinations of symmetrically related binding sites for both anchor and disruptor arm, so as to yield a total of twelve crude, candidate cis-binding geometries.

Second, Claude derived an algorithm which, for each fixed geometry and choice of *k∈{*3, 4, 5, 6, 7, 8*}*, attempts to construct a placement of the alpha carbons of (G_4_S)_*k*_ tandem linker residues so as to yield an unobstructed path between the pair of complementary VHH termini. For the latter, minor intervention on our part was required to ensure that pathological linker geometries (i.e., threading the “donut hole” of the open conformation of IgE C*ε*3–C*ε*4) were excluded, and that certain steric clashes were properly accounted for. The combined procedure then nominates the shortest such *k* for which an unobstructed linker could be constructed, along with the corresponding construction. One such linker construction is provided in Figure 4 (bottom right).

## 4 Results

We nominated designs from both the disruptor and anchor design campaigns for experimental validation and assessed their functional activity via bio-layer interferometry (BLI). For the disruptor arm, we performed a multi-phase, competitive BLI experiment to measure the extent to which the designed antibody in solution can displace IgE bound to an immobilized soluble Fc*ε*RI*α* ectodomain, mimicking the physiological context of Fc*ε*RI on mast cells and basophils. For the anchor arm, we performed standard kinetic characterization of free IgE binding to immobilized VHH.

### 4.1 Designed disruptor variants displace IgE from bound Fc*ε***RI***α*

We first nominated 91 disruptor designs for experimental screening, alongside the parent sdAb026 and single alanine mutants of its C*ε*4-facing CDR3 previously reported to remove functional activity [38], at a single concentration. Twenty designs removed IgE faster than every negative control. We classified these 20 sequences as positive hits for disruptive activity. Six hits appeared to improve upon the activity of sdAb026. Hits originated from every design scope and method, albeit at different rates (Figure 5D). Redesign of CDR2 was by far the most productive route, accounting for most of the hits and most of the designs that appeared to exceed the disruptive activity of the parent. However, activity did not depend on staying close to the parent sequence, and functional activity was retained for sequences differing by as many as eight substitutions (Figure 5B), demonstrating the ability of biomolecular models to navigate sequence functional landscapes zero-shot. Substitutions throughout CDR2 and at two positions in CDR1 appeared in designs more active than the parent (Figure 5E), whereas most positions in the C*ε*4-facing CDR3 never appeared in an active design at all, consistent with the loss of activity in the alanine mutants of that loop.

**Figure 5.**
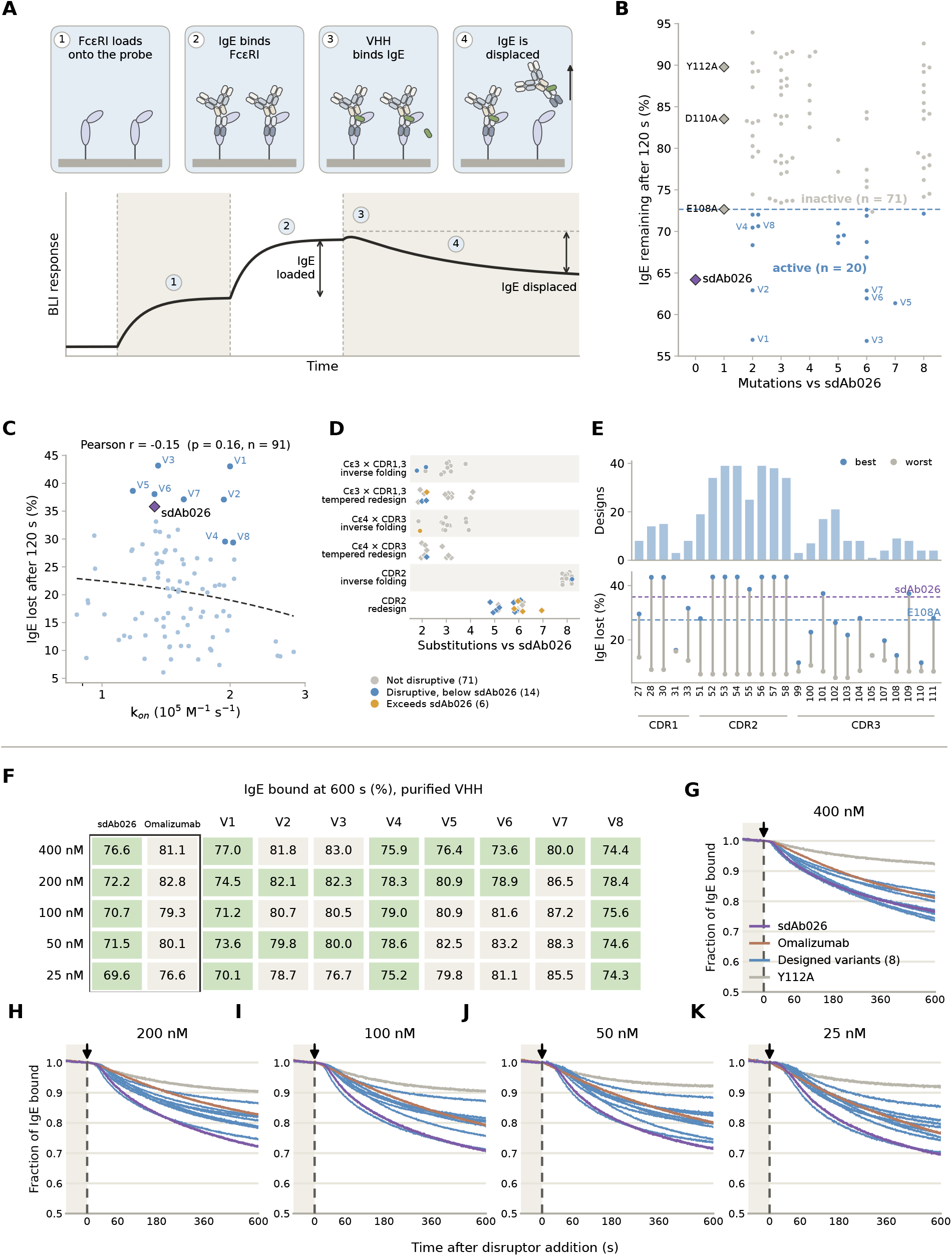
Experimental characterization of disruptor arm designs. **(A)** Schematic and idealized sensorgram from the competitive BLI assay. The IgE displacement is quantified in terms of the decline from the peak BLI response relative to total IgE loading. **(B–E)** Initial screen of 91 designs via a 120 s dissociation phase. **(B)** IgE remaining at the end of the dissociation phase against the number of substitutions from sdAb026. Twenty variants (blue) exhibited an apparent IgE off-rate exceeding every negative control’s; the eight variants carried forward for purification are labeled V1–V8. **(C)** The disruptive effect is not correlated with the association rate in a standard binding kinetics assay. **(D)** The designs by redesign scope and method and by substitution count. **(E)** The number of designs mutated at each CDR position (top) and the best and worst IgE displacement among them (bottom). **(F–K)** Purified variants were assayed at 600 s of dissociation and at five therapeutically relevant concentrations, alongside omalizumab. **(F)** Percentage of IgE bound at 400 to 25 nM; green cells removed more IgE than omalizumab. **(G–K)** Time courses at 400, 200, 100, 50 and 25 nM; the arrow marks addition of the disruptor.

We then purified eight active designs, alongside sdAb026 and the inactive alanine mutant Y112A, and compared them to each other and to omalizumab over a range of therapeutically relevant concentrations (Figure 5F–K). To do so, we performed an extended disruptive BLI assay, with the disruption phase extended to ten minutes. All variants retained disruptive activity greater than the Y112A control at all concentrations. Encouragingly, three variants, V1, V4 and V8, removed more IgE than omalizumab at every concentration tested. However, the variants did not appear to improve upon the parent sdAb026, with the best variant, V1, providing a similar response at most concentrations.

To confirm that disruptive activity is not solely a function of binding, we conducted single-point BLI assays of free IgE binding to immobilized VHH. Nearly all of the designs retained binding activity, many with very slow dissociation constants at the limit of measurement (Appendix E). The apparent association rate, *k*_on_, did not predict disruption activity (Figure 5C). Additionally, the alanine mutant negative controls appeared to bind IgE nearly as tightly as sdAb026, but removed far less IgE in the disruptive BLI screen (Figure 5B). Thus, a single substitution at the C*ε*4 interface (i.e., the negative controls) can abolish disruption while leaving affinity nearly intact.

### 4.2 Epitope-diverse anchor arm hits

We next nominated and experimentally tested 57 anchor arm designs via BLI, with the VHH immobilized on the probe. Three variants exhibited cleanly resolved kinetic fits, for a hit rate of approximately 5% (Figure 6A). The hits derived from two of the three epitope-specific design blocks: the side of C*ε*2 distal to the consensus binding site of variants B1, E1, and F4 [41]; and the C*ε*3–C*ε*4 surface. Somewhat surprisingly, no designs targeting the consensus epitope of variants B1, E1, and F4 exhibited binding. Corroborating recent literature [42], the critic scores do not perfectly anticipate which designs would bind; anchor design 1 was the lowest-ranked of the 19 nominees targeting the distal C*ε*2 surface, whereas anchor designs 2 and 3, were second-best within second of the 15 nominees targeting the C*ε*3–C*ε*4 surface.

**Figure 6.**
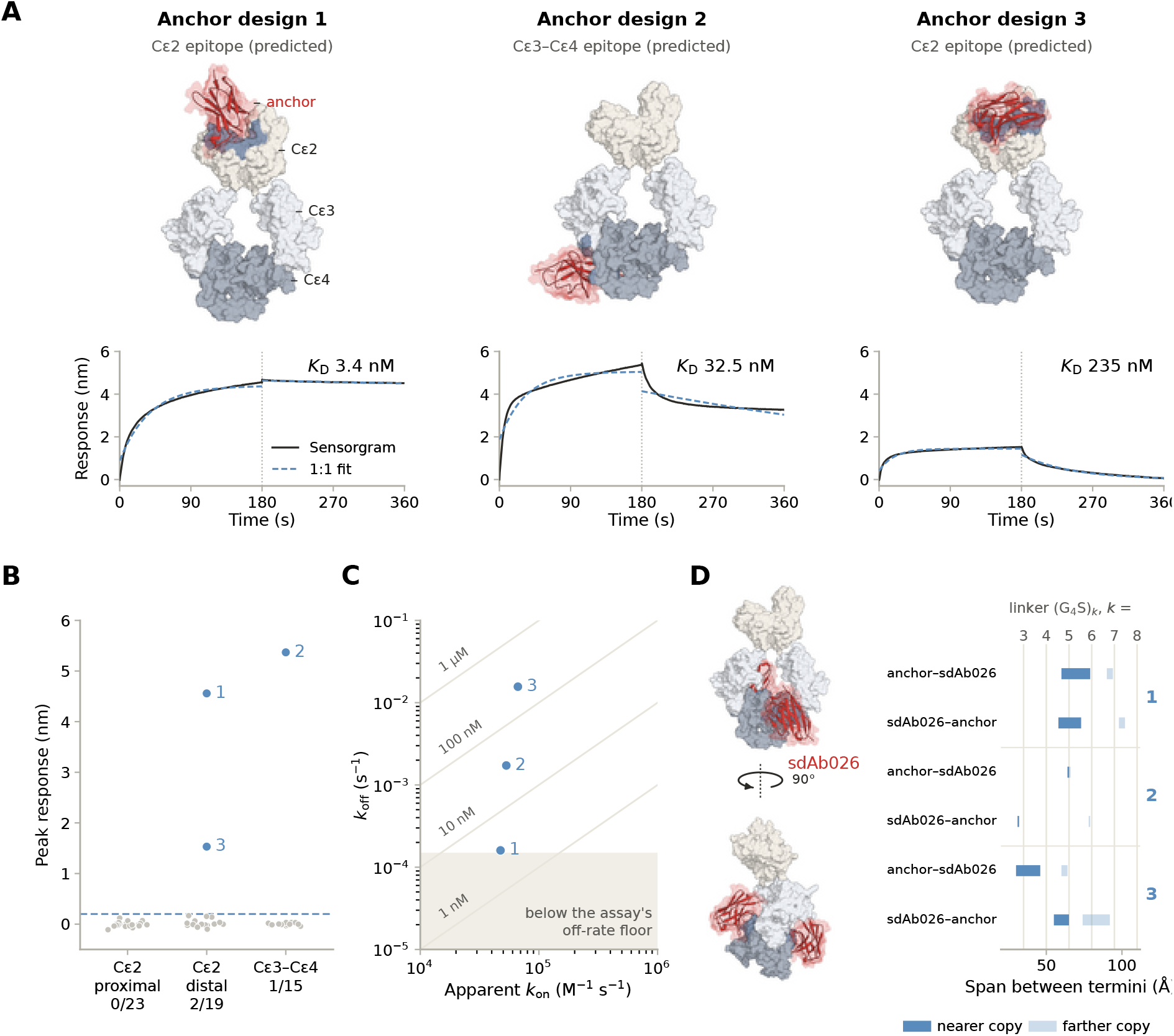
Experimental characterization of *de novo* anchor arm designs. **(A)** Three anchor designs bound IgE, with the predicted pose on IgE-Fc and the BLI sensorgram (black) with a single-exponential fit (dashed) shown. **(B)** Peak association response of all 57 *de novo* anchors by design block; below the dashed line (0.2 nm) no binding was resolved. **(C)** Apparent association and dissociation rates of the three hits, with lines of equal *K*_D_. **(D)** The predicted poses of the anchors suggest possible lengths of (G_4_S)_*k*_ linkers that would combine with sdAb026 into a tandem VHH biparatopic construct, in either orientation. Distance ranges are derived from the two C*ε*2 conformations.

Plotting the association rate against the off-rate (Figure 6C) reveals the three hits associate at similar rates, so their range of apparent affinities, from 3.4 to 235 nM, arises almost entirely from their off-rates. We caveat, however, that these affinity measurements are potentially confounded by avidity; because IgE presents each Fc domain twice, binding events to adjacent immobilized VHHs may not be independent. Curiously, though, anchor designs 1 and 3 appear to occupy a near-symmetric, single binding site at the intersection of the two IgE monomers (Figure 6A). Experimental characterization of avidity will thus provide an unusual mechanism by which to directly probe the accuracy of the predicted binding pose.

Using the predicted binding poses, we explored the linker lengths required to join the anchors with the original disruptor, sdAb026, when the latter is bound to the same IgE dimer (Figure 6D). Because there are two identical binding sites for sdAb026 and two linking orders, there are four possible orientations of the two arms. For all three anchors, at least one orientation admits a linker length within distribution for 3–5 (G_4_S) repeats, a typical range for therapeutic antibodies. Anchors 2 and 3 are closer to the sdAb026 binding site, and hence appear to be more compatible with one tandem VHH orientation over the other. Because these distances have some flexibility depending on the conformation of C*ε*2, we did not commit to a single linker, but built a library (using our disruptor variants in place of sdAb026) that covers both orientations and a range of linker lengths for each anchor (Section 3.4) for further characterization.

## 5 Discussion and future work

In this report, we demonstrated that computational design can produce two functional arms of a biparatopic as the first key step towards a best-in-class IgE disruptive inhibitor. Our parallel design campaigns produced (a) a disruptor arm that removes IgE from Fc*ε*RI, and (b) an anchor arm that binds to IgE at a noncompetitive epitope. Through Claude agentic orchestration of biomolecular generative models [42], both were obtained within weeks rather than the months expected of traditional screening workflows.

Starting from sdAb026, a single-domain antibody with previously reported disruptive activity [38], we obtained 20 variants with measurable disruptive activity. These variants were up to eight substitutions from the parent and represent a diverse functional family different from one another by as many as ten residues (out of 31 CDR residues). These findings mirror similar results in the use of biomolecular foundation models to design functional sequences far from original starting points [50–52]. We will continue to explore agent-orchestrated computational design campaigns [42] to venture further from known sequence space, steering towards other properties of interest in addition to preserving function. While we did not measure any variants with improved functional activity over sdAb026 at controlled doses, diversification of existing functional sequences provides multiple starting points for independent optimization for properties beyond functional potency, such as developability, biparatopic compatibility, and cross-reactivity. This represents a significant reduction in time and cost over the custom library-scale screening protocols previously required for such diversification [32, 33].

In combination with our anchor designs, we expect biparatopic construction (as tandem VHH or as VHH-Fc) to provide a significant boost to potency and disruptive efficacy. Indeed, the parental disruptor sdAb026 was reported to exhibit disruption at therapeutically relevant concentrations only when combined with an anchoring VHH [35, 37, 41]. Similarly, the orthosteric disruptor DARPin E2_79 experienced dramatically improved potency after conversion to a biparatopic format [27, 39]. In this light, we are encouraged to observe improved activity over omalizumab even when our designs are expressed as monovalent constructs.

A key limitation of the disruptor design methodology pursued here is the reliance on an existing sequence (and in fact, docked structure) of a molecular species with the desired functional activity. Future work will explore if Claude can employ reasoning strategies to design functional sequences when no reference sequence is available; or even more ambitiously, when the structural specification of the desired function (in this case, inducing disruption by biasing IgE Fc toward a closed conformation) is entirely unavailable.

The subcomponents generated here by Claude will provide an excellent basis for the development of candidate biparatopics for further characterization towards a best-in-class IgE disruptive inhibitor. Following biophysical validation of free and bound IgE inhibition, candidates will be tested in basophil and mast-cell assays for disruption, activation, and protection against allergen challenge across donors, followed by further lead optimization and functional, pharmacokinetic, and pharmacodynamic studies in animals. A key risk will be to avoid anaphylactogenicity via receptor cross-linking; we anticipate that geometric reasoning and analysis of biparatopic formats with Claude will prove highly useful in this next stage of discovery.

Looking beyond IgE, the strategies and methods developed in this work create a powerful foundation for additional campaigns of rationally designed antibodies capable of more complex biological functions, opening the door for best-in-class medicines across human diseases.

## Acknowledgements

We thank Adu Subramanian, Nathan Mallipeddi, and Dan Levin for their early feedback on our thesis; Nathan Robertson, Pascaline Mary, David Holmes, Assel Ismoldayeva, Sam Sledzieski, Jack Kenney, and the entire Divergence Labs team of determined enthusiasts for their experimental guidance and program support; Shahram Salek, Leonard Presta, and Joao Vieira especially for their invaluable expertise and strategic guidance on this program as well as the entire Divergence Labs R&D Committee and Board of Directors for their support.

## A Technical details of disruptor redesign

We provide extended commentary and technical details for all disruptor redesign phases as outlined in Figure 3.

### A.1 Characterization of 5NQW interface geometry and clarification of redesign scopes

#### Determination of design scope

After analyzing the binding mode of sdAb026 in PDB 5NQW, Claude nominated the C*ε*3 interface, C*ε*4 interface, and free scopes detailed in Figure 7. As reported in the main text, the free scope was eventually dropped in favor of a fourth, interface-distal CDR2 scope so as to increase the diversity and extent of the explored optimization space. We emphasize that chronologically, full redesign was initially carried out against the three non-CDR2 scopes before the conceptualization of the fourth, complementary CDR2 scope reported in Figure 3. Claude then further identified the first three, interface-proximal scopes as being candidates for targeted mutagenesis (edit distance 2-4) rather than full scope redesign, utilizing both inverse folding (Appendix A.2) and a gradient-based mutagenic design scheme developed for precisely this purpose, as described in Appendix A.3.

**Figure 7.**
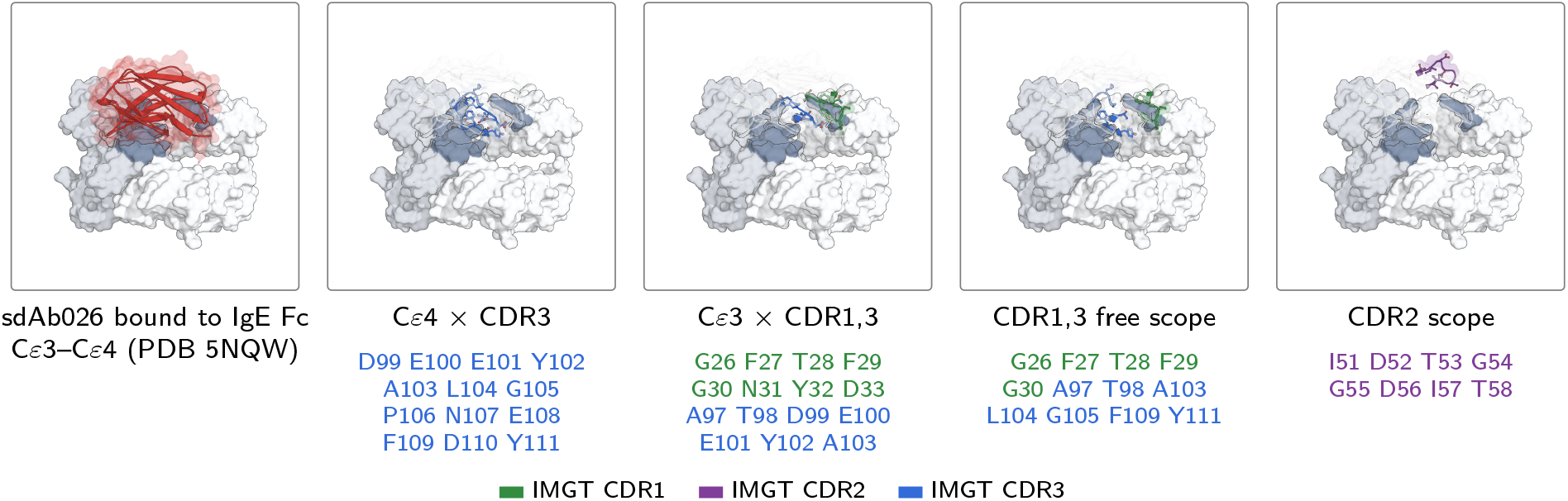
Redesign scopes on sdAb026. sdAb026 bound to IgE Fc C*ε*3–C*ε*4 (PDB 5NQW; C*ε*3 and C*ε*4 rendered in white and light-grey, respectively). Left-middle to right: the four redesign scopes with color-coded scope residues highlighted according to their respective CDRs.

### A.2 Inverse Folding of Scopes 1-3 Sequence-biased inverse folding

To achieve the desired mutagenic effect with inverse folding, Claude initially considered combining a uniform and calibrated sequence bias with SolMPNN [46]. First, *N* = 250 unbiased draws using SolMPNN at *T* = 0.1 were produced for each scope *S*, with 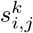 denoting the *k*-th recorded logit of the *j*-th residue member of the *i*-th draw, after which Claude sought to calibrate a uniform, scope-specific biasing parameter *b*_*S*_ so as to maximize the fraction.

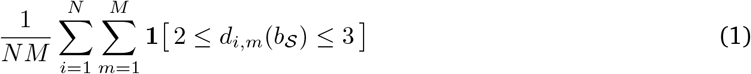

where

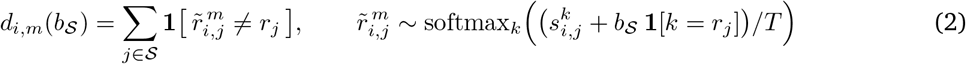

of biased resamples predicted to land between two and three edits, where *r*_*j*_ is the parent sdAb026 residue at scope position *j*, 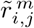 is the *m*-th of *M* residues resampled at that position from the recorded logits 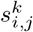 of unbiased draw *i* after adding the bias *b*_*S*_ to the parent residue’s logit (softmax over the twenty amino acids *k*, tempered by *T* ), *d*_*i,m*_(*b*_*S*_) is the number of substitutions within *S* carried by that resample, and **1**[*·*] is the indicator function. Claude performed this calibration procedure to obtain such a *b*_*S*_ for each of the three interface-proximal scopes. Logo diagrams for the averaged unbiased and biased position-specific probability matrices are shown on the top and bottom, respectively, of Figure 8 (b), emphasizing that bias

**Figure 8.**
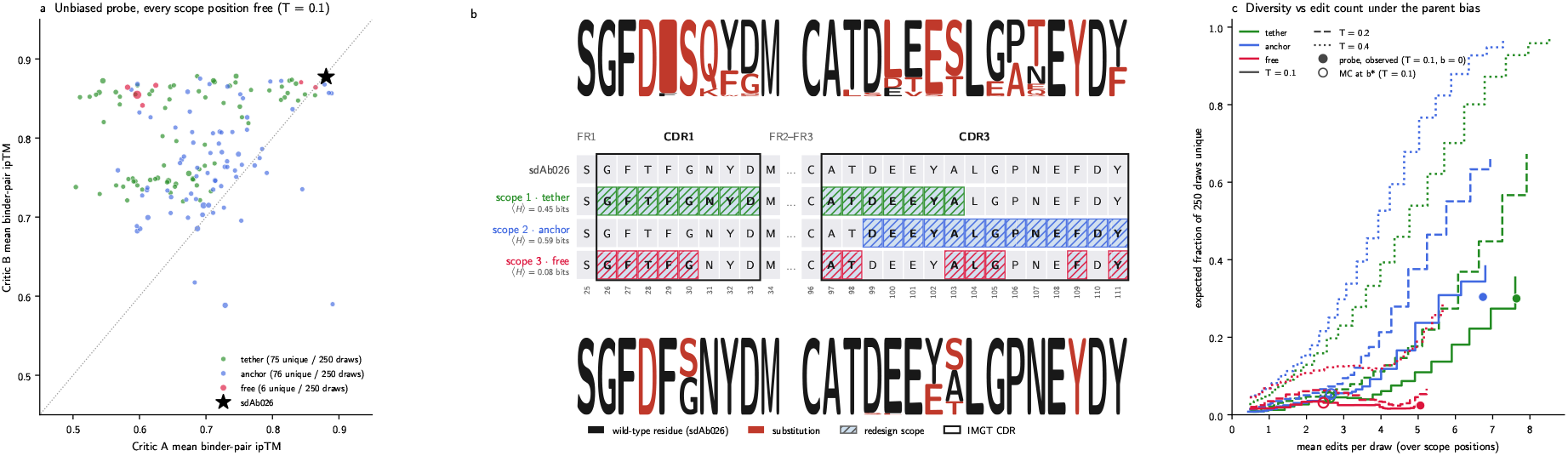
Sequence-biased inverse folding of scopes 1–3. (a) Critic A against Critic B mean binder-pair ipTM for the unique sequences among the *N* = 250 unbiased SolMPNN draws per scope at *T* = 0.1, with marker size scales with the number of draws yielding the sequence. (b) Sequence logos of the unbiased draws (top) and of the biased resamples of Equation (1) at the calibrated *b*_*S*_ (bottom), over sdAb026 with the three scopes marked; black, the sdAb026 residue; red, a substitution. (c) Expected fraction of unique sequences among 250 biased resamples against their mean edit distance as *b*_*S*_ is swept, at *T* = 0.1, 0.2 and 0.4; filled markers, the unbiased draws.

was primarily applied toward a small number of higher entropy indices, so as to yield an undesirable tension between edit distance and redesign diversity, and so that the realization of diverse sequences requires a stronger bias, yielding designs of both high edit distance and low unbiased model likelihood. Indeed, as modeled in Figure 8 (c), the fraction of unique designs rises only modestly within the edit distance interval of interest (2-4 edits) as temperature is increased from *T* = 0.1 to *T* = 0.4. Subsequent inverse folding and critic evaluation substantiated such biasing as detrimental to model confidence, suggesting that biasing in this fashion yields sequences which are consistent with neither the inverse folding model itself nor the downstream folding critics.

#### Inverse Folding with Manually Constructed Scopes

In light of poor critic confidence on the part of manually calibrated biased inverse folding, Claude elected instead to manually enumerate all double-, triple-, and a select number of quadruple mutants of sdAb026 until a predetermined budget of *N* = 750 was obtained, and subsequently inverse fold at *T* = 0.4 with SolMPNN. Critic confidence with respect to both nominated critics, edit distance, and DockQ are visualized in Figure 10. In particular, we report both mean critic *binder-pair* ipTM (that is, the average ipTM between the design and the respective IgE dimer chains), and the *sub-interface DockQ mean* [53, 54], which we define as the average DockQ of the variant with respect to both C*ε*3 and C*ε*4 interfaces.

### A.3 Tempered redesign of Scopes 1-3

In addition to the inverse folding in Appendix A.2, Claude enabled the rapid development and integration of a second, gradient-based design approach we’ve termed **tempered redesign**, which together with an improved **interface geometry loss**, yields designs which achieve both higher critic confidence and deviate less significantly from the intended reference geometry across all three initial design scopes, and as visualized in Figure 9. A more extensive technical discussion of both techniques can be found in Appendix C.

**Figure 9.**
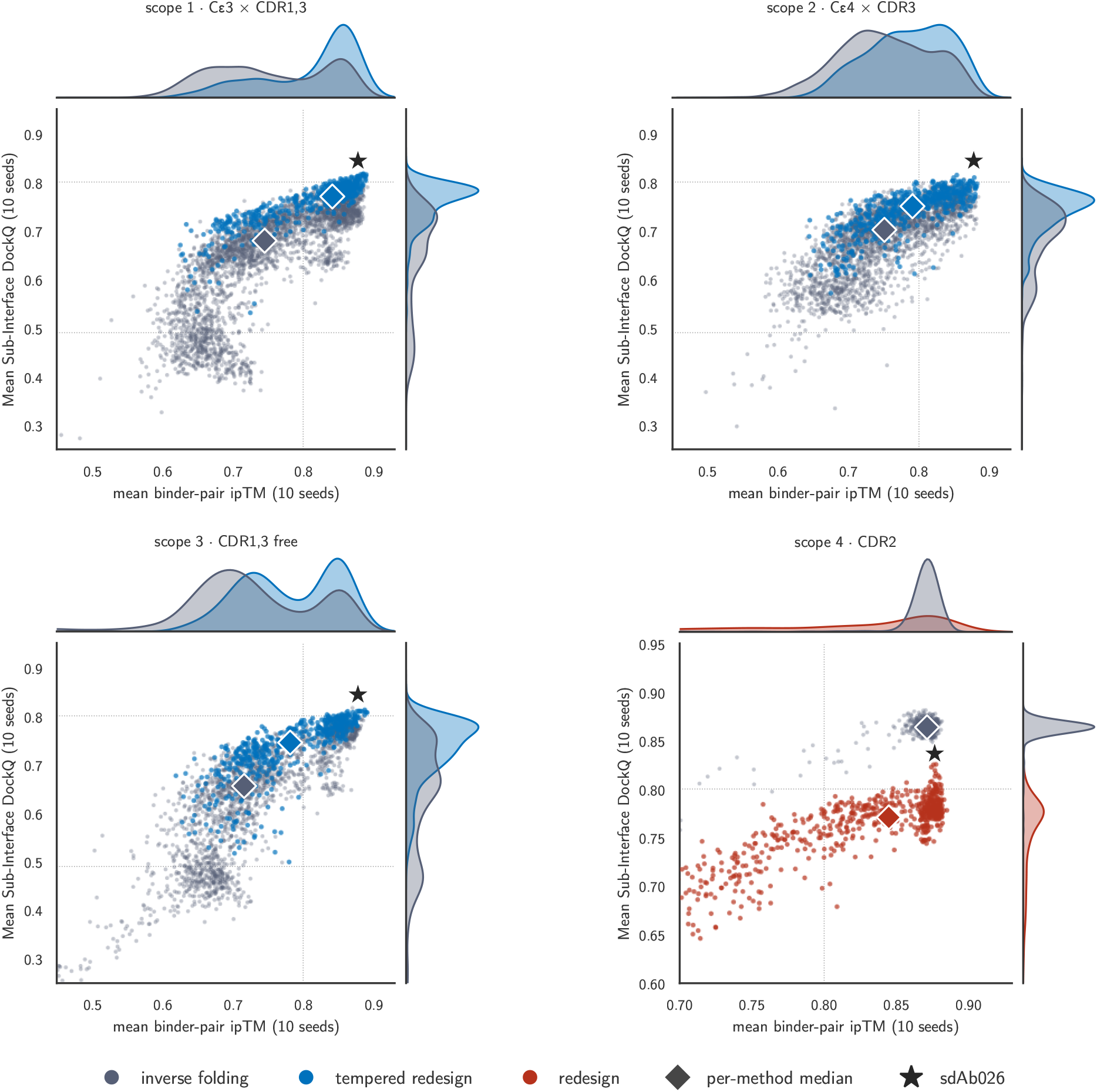
Summary of sdAb026 redesign designs. Top-left, top-right, and bottom-left: scatters of mean binder-pair ipTM against mean sub-interface DockQ for double, triple, and quadruple mutants generated by tempered redesign (blue) and inverse folding (grey) against the C*ε*3 interface, C*ε*4 interface, and free scopes, respectively. Bottom-right: scatter of mean binder-pair ipTM against mean sub-interface DockQ for the redesigns obtained via inverse folding with SolMPNN (grey) and gradient-based redesign (red) against the CDR2 scope.

**Figure 10.**
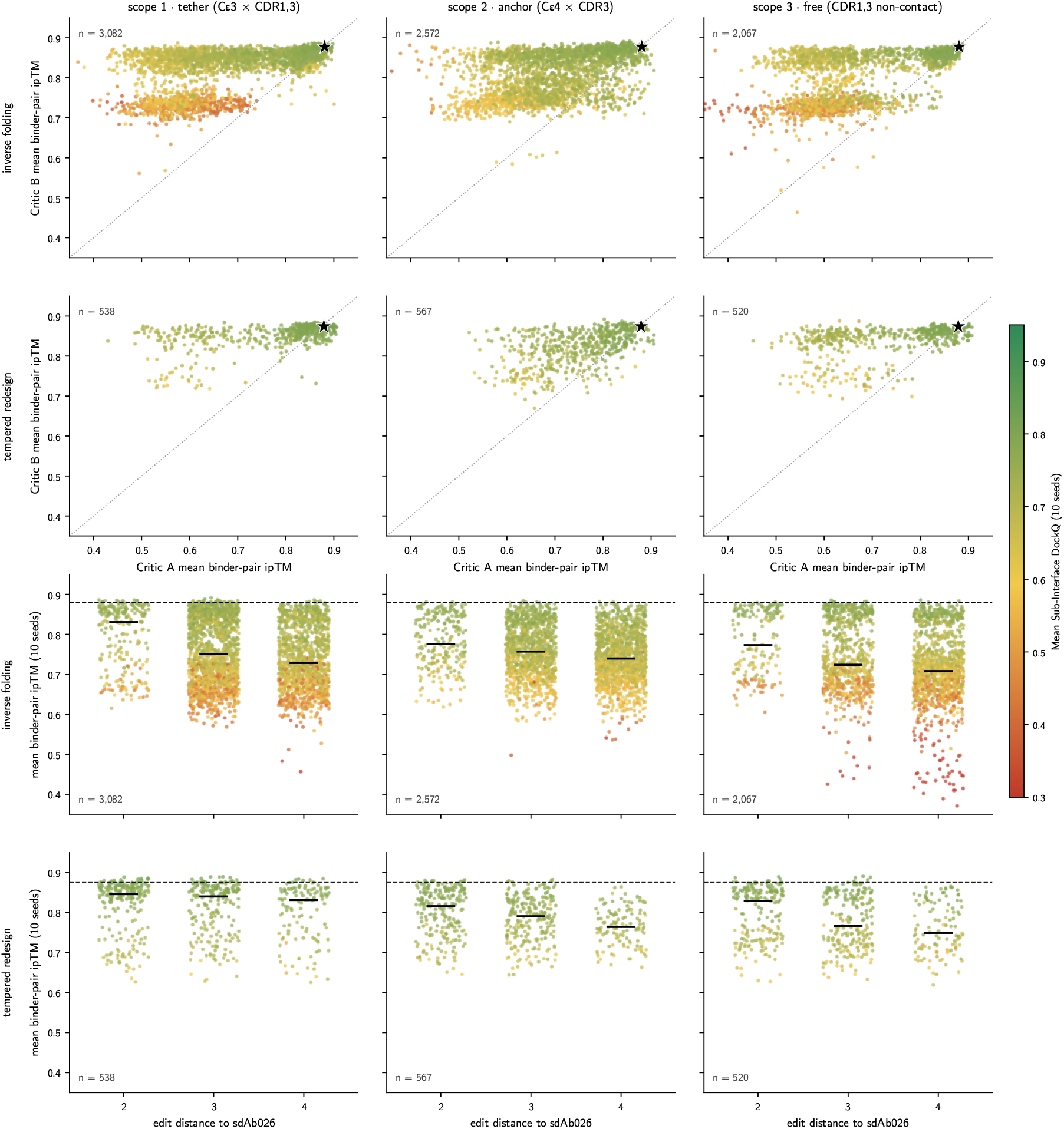
Inverse folding and tempered redesign of scopes 1–3. Rows: inverse folding (every unique sequence among the enumerated double, triple and sampled quadruple mutants of sdAb026) and tempered redesign (the production pool at two to four substitutions); columns: the three scopes. Top: Critic A against Critic B mean binder-pair ipTM per sequence, colored by mean sub-interface DockQ to PDB 5NQW over ten seeds; star, sdAb026. Bottom: edit distance to sdAb026 against mean binder-pair ipTM over ten seeds, same coloring; bars, per-edit medians; dashed line, sdAb026.

### A.4 CDR2 Redesign

For the CDR2 scope, and due to its distance from both C*ε*3 and C*ε*4 interfaces, Claude near-autonomously executed full-scope redesign using a combination of both inverse folding with SolMPNN and gradient-based redesign. Toward this end, SolMPNN (v_48_020) was used to generate 750 redesigns at each rung of the temperature ladder *T∈ {*0.1, 0.3, 0.5, 0.7, 1.0*}*, obtaining unique-sequence fractions of 18.9%, 61.6%, 87.6%, 97.9% and 100% respectively (750 draws per rung), of which 1,289 unique designs over the ladder were scored under both critics (Table 1). Gradient-based redesign was similarly run with identical interface geometry loss settings to those developed and used in Appendix A.3 to obtain 750 trajectories, yielding 695 distinct designs (93%). The results of both inverse folding and gradient-based redesign are displayed in Figure 9, indicating that direct redesign trades off exact recapitulation of interface geometry as measured by mean sub-interface DockQ for an increase in mean critic confidence in the form of a thicker tail.

### A.5 Conclusion and nomination

Claude concluded the round by nominating a total of 91 distinct designs by selecting the top 13 from each of four total C*ε*3 and C*ε*4 scope-method combinations, as well as 20-each from each of the two design approaches employed against the CDR2 scope. One of the CDR2 scope variants redesigned using inverse folding was subsequently dropped to accommodate an additional control, yielding the reported total of 91 designs. An overview of the ipTM of the nominated designs and their edit distances to sdAb026 is provided in Figure 4. Additional design details are provided in Table 1.

## B Technical details of de novo anchor design

### B.1 Computational Prediction of Literature-Described C*ε*2 Binders

In Figure 11 (a), we list reported affinities and kinetic quantities for immunization-derived, non-humanized nanobodies confirmed to bind IgE Fc [41]. We additionally pursue refolding against both IgE Fc C*ε*2 and C*ε*2–C*ε*4 dimers using one of the two nominated campaign critic models and report intra-variant epitope Jaccard and ipTM results in Figure 11 (b, c). In particular, we obtain a shared, predicted consensus epitope for variants B1, E1, and F4, but fail to reach consensus for variants B3 and D2. Folds against C*ε*2–C*ε*4 and C*ε*2-only dimers are provided in Figure 11 (d).

**Figure 11.**
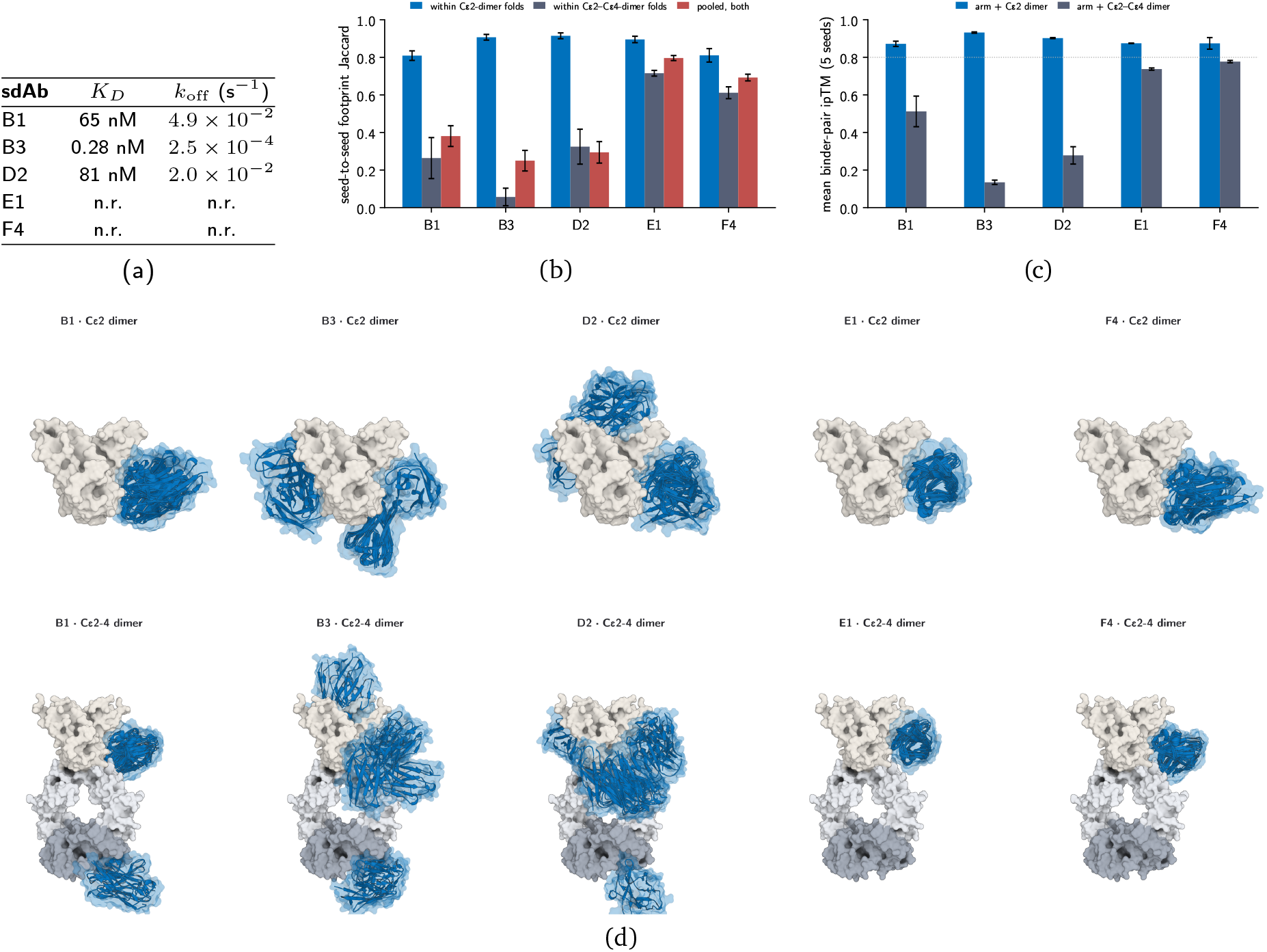
Literature-described C*ε*2 anchors: reported kinetics and their refolding on the C*ε*2 dimer and on the C*ε*2–C*ε*4 dimer. (a) Reported BLI affinity and dissociation rate on IgE-Fc for five experimentally characterized sdAbs from Spillner et al. [41] (n.r.: not reported.) (b) Seed-to-seed consistency of the 4 Å footprint as the pairwise Jaccard within the five C*ε*2-dimer folds, within the five C*ε*2–C*ε*4-dimer folds, and pooled over all ten (mean *±*SEM over pairs; Critic A). (c) Mean binder-pair ipTM *±*SEM over the five Critic A seeds on each target. (d) All ten refolds of each sdAb (Critic A and Critic B, five seeds each) superimposed on one reference surface of the target, with sdAb chains rendered in blue.

### B.2 Computational design against C*ε*2 Scope

#### Scope construction

We consider a global C*ε*2 scope constructed from PDB 2Y7Q as the set of solvent-accessible residues (per-residue SASA*≥* 15^2^ via PyMOL) whose alpha carbon has an unobstructed line of sight of length at least 20 Å to an alpha carbon on C*ε*3–C*ε*4. We then further restrict to obtain a local and limited *B1-consensus scope* as the intersection of this initially constructed C*ε*2 scope with the set of residues present in the epitope footprint in at least 75% of B1 cofold predictions from Appendix B.1. Both scopes are visualized in Figure 12. We next describe a total of three Claude-driven de novo design phases against these two scopes.

**Figure 12.**
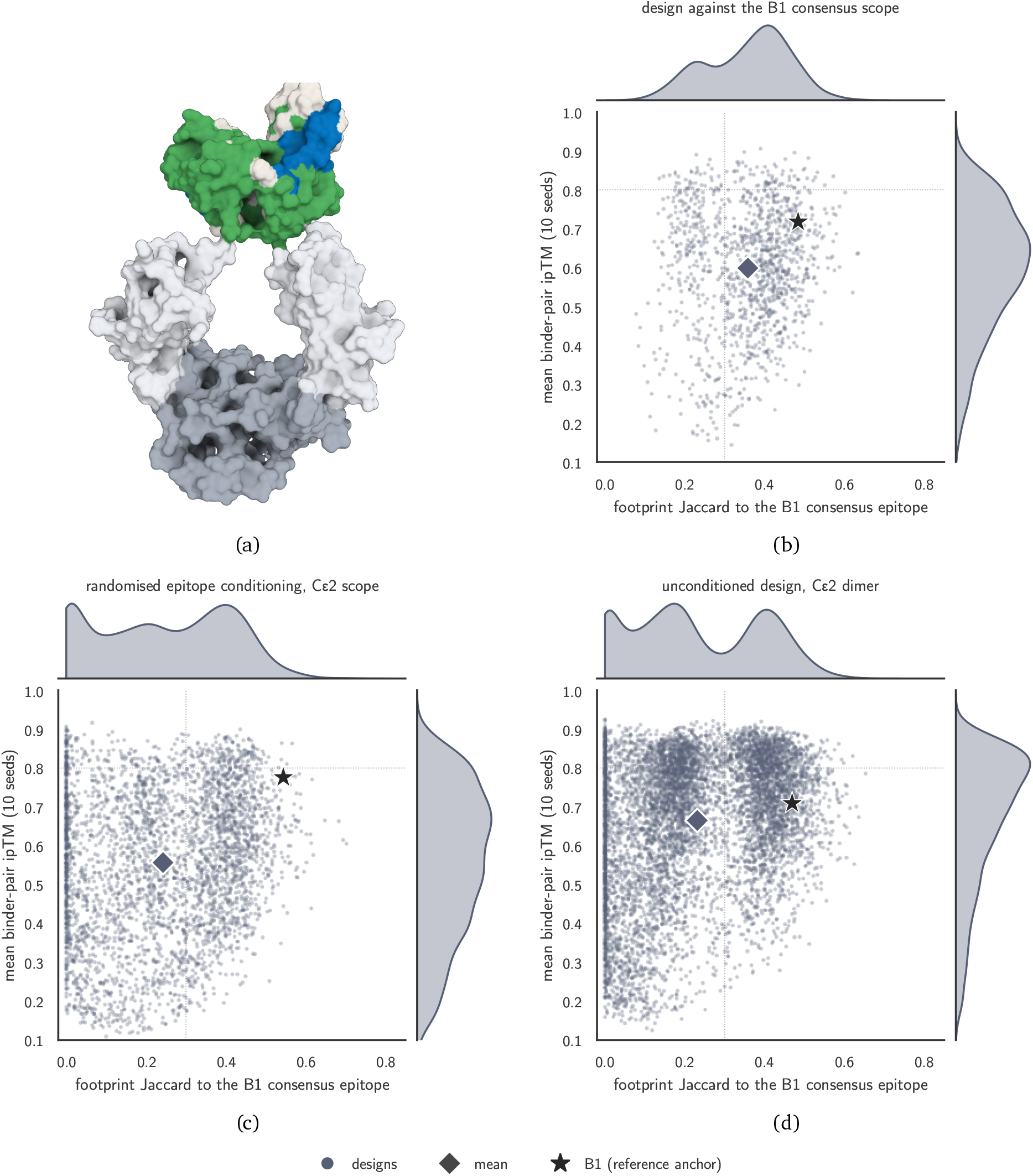
The C*ε*2 design scope and the three design rounds against it. (a) The C*ε*2 inside-face scope (green) and the nine-residue B1 consensus epitope (blue) on the receptor-bound IgE Fc of PDB 7SHT, and with all domains rendered as surfaces. (b–d) Mean binder-pair ipTM over the ten critic seeds against the footprint Jaccard to the B1 consensus epitope (mean over the Critic A seeds), one point per unique design, with marginals: (b) design against the B1 consensus scope, (c) epitope scanning over the C*ε*2 scope, (d) unconditioned design against the C*ε*2 dimer.

#### Design against B1 consensus scope

For each of the twelve distinct frameworks in Table 2, Claude ran 102 design trajectories of epitope-conditioned redesign of all three CDRs, for a total of 1224 total design trajectories. Each such design was then subsequently refolded using five distinct seeds and across both campaign critic models, with results reported in Figure 12. We then iteratively refine the top-40 variants identified by 10-seed mean ipTM through repeated inverse folding and folding, similar to the procedures described by Fry et al. [47], Frank et al. [48], Cho et al. [49]. Results are reported in Figure 12, including both 10-seed mean critic confidence and epitope Jaccard to the target epitope.

#### Epitope scanning against C*ε*2

To increase epitope diversity, Claude subsequently elected to perform design with randomized epitope selection: a query residue is randomly selected from the global scope from which an epitope is then constructed as the set of all in-scope residues whose alpha carbon lies within a 12 Åcutoff of the query. We henceforth refer to this approach as *epitope scanning*. Design is then carried out as in the previous phase, and against the C*ε*2 dimer, with the additional repulsive loss term applied to out-of-scope residues on C*ε*2. In total, this round generated a total of 3600 distinct trajectories evenly divided among the same twelve frameworks shown in Table 2. The top 75 designs by 10-seed mean ipTM were subsequently nominated for further iterative refinement totaling 50 additional H100 hours, and yielding an average of 20 cycles of refinement. Results are reported in Figure 12, including both 10-seed mean critic confidence and epitope Jaccard to the epitope from the previous round.

#### Unconditioned design against C*ε*2

Finally, Claude elected to run a final, unconditioned design run against the C*ε*2 scope, this time without any repulsive loss term. This round generated a total of 7194 design variants across the 12 VHH frameworks. Further experimental details can be found in Table 3, and results are reported in Figure 12.

### B.3 Computational design against C*ε***3–C***ε*4

#### Linker- and epitope-related filtering procedure

The flexible positioning of IgE C*ε*2 relative to C*ε*3–C*ε*4 makes it hard to gauge the geometric compatibility of an anchor placed on the latter with a disruptor on the former on tandem or Fc-fusion linker geometry, as doing so would require more precise modeling of domain rearrangement and linker geometry to accommodate cis binding of the resulting biparatopic. In contrast, the comparatively rigid C*ε*3–C*ε*4 varies largely between the open and closed conformations described in Section 2, so that direct screening for linker compatibility is more straightforward. For this reason, Claude formulated a screening algorithm for determining the compatibility of a choice of anchor with a C-terminal IgG1 fusion, as in [41], and together with disruptor sdAb026 (via PDB 5NQW). The procedure involves first performing a rigid alignment of both the anchor arm as well as one of the two symmetrically related sdAb026 chains in 5NQW to PDB 2Y7Q. Then, for each of linker lengths (G_4_S)_1_ (*n* = 4) and (G_4_S)_2_ (*n* = 8), we construct circular shells centered at the respective VHH N-termini, and with inner and outer radii equal to 0.8 *×L* and 1.2 *×L* for *L* ≜ 0.8 *×*3.8 *×n*. A 1 Åcircular grid is then constructed about the midpoint of the N-termini, and a grid search is performed for apices which satisfy strict clash criteria. Namely, the shortest clash-free paths are identified via Dijkstra’s shortest-path algorithm on a 1.5Å cubic lattice, with lattice points within 3 Åof IgE Fc, receptor, or arm chains excluded. A final check is applied by screening for clashes involving a rigid-body cylindrical approximation of the IgG1 Fc. If such a non-clashing path exists, the anchor is said to be linker-compatible. We additionally apply a secondary, epitope-related filter removing any design which overlaps with the respective residue footprints of sdAb026 or Fc*ε*RI*α*.

#### Epitope scanning against C*ε***3–C***ε*4

In line with *de novo* design against C*ε*2, Claude similarly pursued design against C*ε*3–C*ε*4 with an identical epitope scanning procedure to the one described in Appendix B.2, and with a similar repulsive avoidance loss intended to exclude residues within the footprints of sdAb026 and Fc*ε*RI*α*. A total of 1584 design trajectories were run, after which the top-50 designs by mean critic ipTM passing both linker compatibility and epitope filters were selected for further refinement in the same manner as described in the C*ε*2 epitope scanning phase, significantly diversifying the set of high-confidence, filter-passing designs, and as shown in Figure 13 (top-right, middle-left).

**Figure 13.**
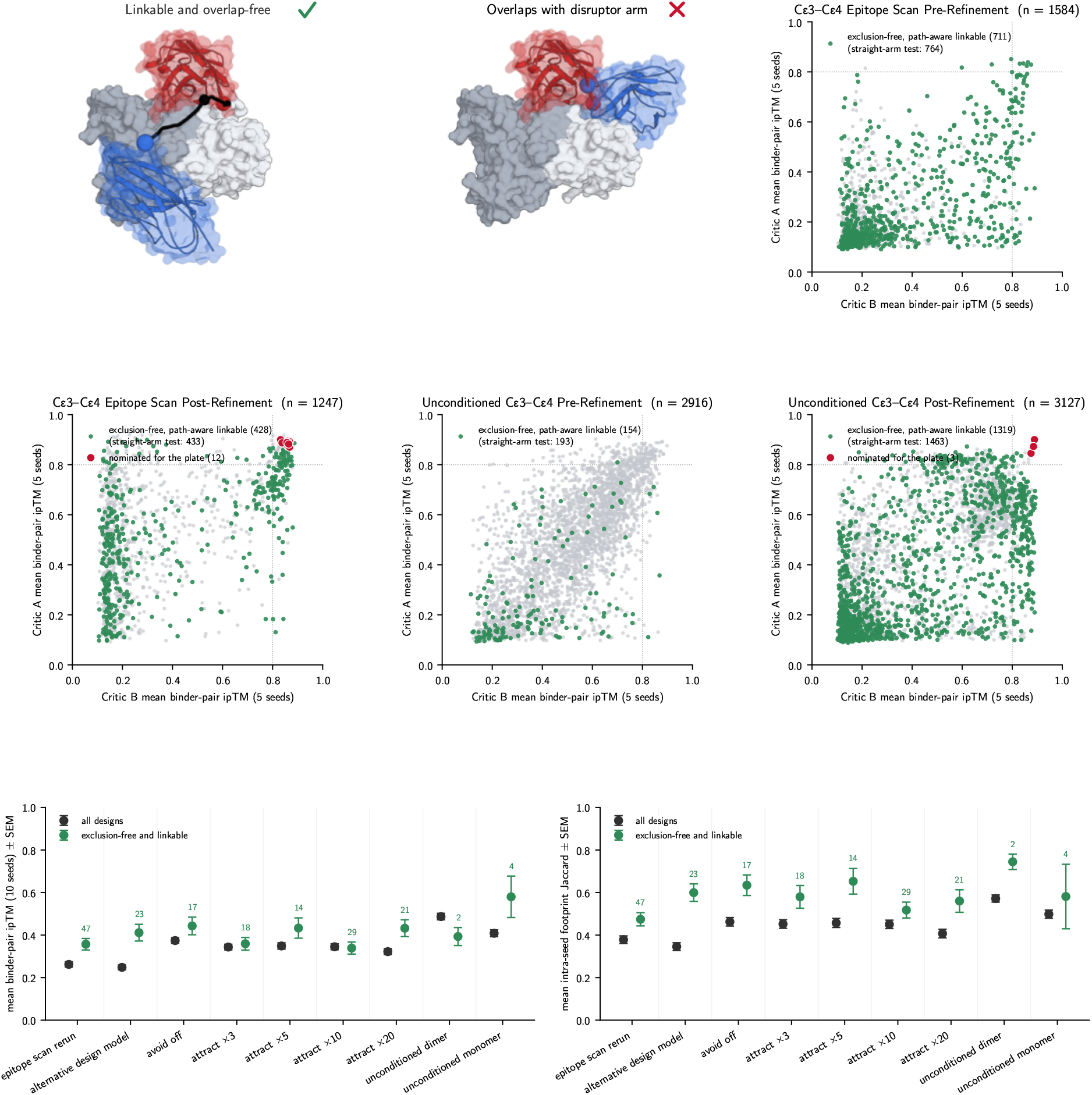
De novo anchor design against C*ε*3–C*ε*4. Top: a nominated anchor passing both linkability and epitope-overlap criteria, tethered to the proxy disruptor arm sdAb026 (red) by a (G4S)_1_ linker proxy (black; arms drawn as shortest obstacle-free paths as described in Appendix B.3) on the 5NQW Fc C*ε*3– C*ε*4; an anchor with epitope (and steric) overlap with sdAb026; the epitope-scan pool before refinement, Critic A vs Critic B mean binder-pair ipTM (5 seeds), with exclusion-free and KIH-linkable designs in green. Middle: the same pool after refinement (plate nominees in red), and the unconditioned pool before and after refinement (plate nominees in red). Bottom: mean 10-seed binder-pair ipTM and mean intra-seed footprint Jaccard, both*±* SEM and per ablation setting, with all designs (black) against the exclusion-free and linkable subset (green; subset sizes above the points).

**Figure 14.**
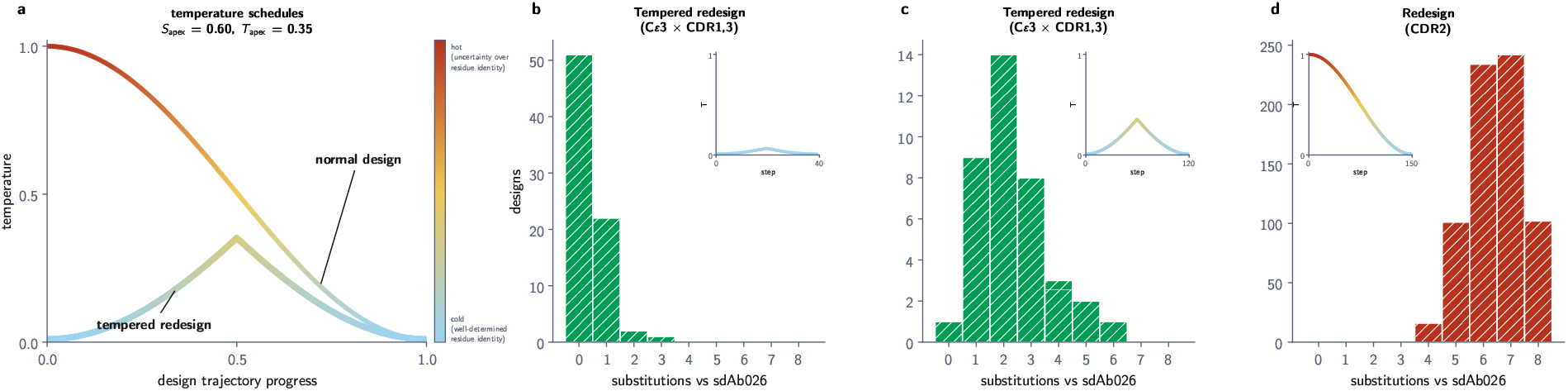
Tempered redesign. (a) Temperature schedules: normal design follows the cosine schedule from hot to cold; tempered redesign runs the same schedule from cold up to the turnaround *S*_apex_ in reference time and back, drawn at *S*_apex_ = 0.60 (*T*_apex_ = 0.35). (b) Substitutions per design in the ultra-low-edit sweep cell (C*ε*3*×* CDR1,3 scope, *S*_apex_ = 0.85, *N* = 40 steps, 76 trajectories). (c) The chosen production cell (C*ε*3 *×*CDR1,3 scope, *S*_apex_ = 0.60, *N* = 120 steps, 38 trajectories). (d) Normal design of the CDR2 scope under the standard schedule (*N* = 150 steps, 695 designs). Insets depict the respective temperature schedules.

**Figure 15.**
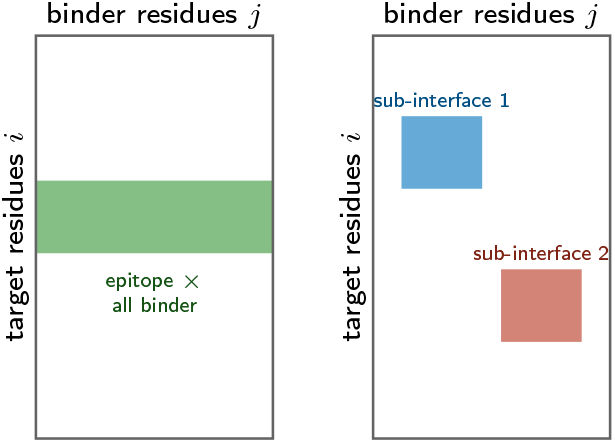
Left: vanilla epitope loss from Candido et al. [55] promotes contact between a specified epitope and the entire binder. Right: the granular interface geometry loss promotes contact between *multiple* distinct epitopes, each with respective, distinct paratope groups not equal to the entire binder.

#### Design ablations against C*ε*3–C*ε*4

The low incidence of filter-passing, high-confidence designs prior to refinement in the previous phase prompted Claude to suggest a series of ablations applied to the design procedure, swapping the design model checkpoint, dropping the avoidance weight, varying the weight applied to the interface geometry loss term, dropping conditioning altogether, and designing instead against a C*ε*3–C*ε*4 monomer. For each, 150 designs were run with the ablated settings, with results reported in Figure 13 (bottom row). Overall, and among other observations, we find that removing the repulsion loss raises the average design confidence as well as the average intra-design epitope Jaccard score above the baseline, albeit at the expense of a reduced incidence of filter-passing designs.

#### Unconditioned generation against C*ε***3–C***ε*4

Citing a desire for consistency with the C*ε*2 design campaign, higher intra-design epitope consistency, and the possibility of hit-expansion as a means to mitigate initially low filter-passing rates, Claude elected to carry out a final round of completely unconditioned design against C*ε*3–C*ε*4 totaling 2916 design trajectories. Of these, a small subset (*n* = 8) was advanced to hit-expansion and diversification via inverse folding with SolMPNN, yielding a refined pool of over 3000 additional designs. Pre- and post-refinement results are shown in Figure 13.

### B.4 Conclusion and nomination

Claude finally nominated a total of 57 designs for experimental validation, including four designed against the B1 consensus epitope on C*ε*2, 38 from subsequent design against C*ε*2, and 15 against C*ε*3–C*ε*4. Further design and model details can be found in Table 3.

## C Claude-driven methodological extensions

Throughout the campaign, we repeatedly leaned on Claude to enable rapid, on-the-fly conceptualization, implementation, and deployment of methodological extensions of open-source and internal design work-flows custom-tailored to accommodate structural, sequential, or other experimental priors specific to the campaign. Two such examples came up during disruptor redesign which we believe materially shifted our capacity to prosecute the campaign, and which we expect to be of further use within our internal design platform going forward. We describe each now as follows.

### C.1 Targeted mutagenesis via tempered redesign

Our design platform consists of a gradient-based design model similar to that disclosed in Candido et al. [55], and which natively supports the redesign of particular residue subsets *starting from scratch*. In conjunction with Claude, we reasoned that by modifying the annealed temperature schedule to instead start at a low, near-zero temperature before rising to a predetermined apex, and then cooling back down to the starting temperature, we could realize a gradient-based mutagenic design scheme. In particular, recall the cosine temperature schedule

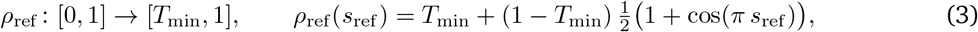

so that the tempered redesign schedule *ρ*_tempered_ is realized as the composition *ρ*_ref_ *π*_transform_, with the tempering transform *π*_transform_ defined as

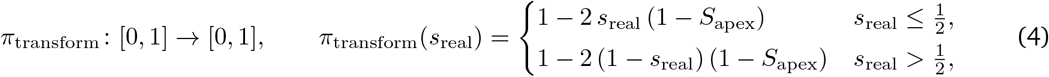

Here, *T*_min_ denotes the minimum design temperature, and *S*_apex_ denotes apex reference time. **Tempered redesign** is then achieved by taking the initial sequence chosen explicitly rather than randomly and applying Algorithm 11 of Candido et al. [55] with tempered temperature schedule *ρ*_tempered_ in place of *ρ*_ref_ .

### C.2 An improved interface geometry loss

An additional design modification whose rapid campaign integration was driven by Claude came in the form of a generalization of an existing epitope loss from Candido et al. [55], allowing for the specification of interface geometry at arbitrary granularity. This **interface geometry loss** enables the creation of multiple distinct interface sub-groups, and the specification of group-specific epitope and paratope residue sets each with distinct distance cutoffs and contact parameters. We provide an overview of this modified interface geometry loss in Figure 16 as well as a demonstration as to the degree to which it can, in combination with tempered redesign, yield redesigns which more faithfully recapitulate the desired interface geometry when compared to the vanilla epitope loss.

**Figure 16.**
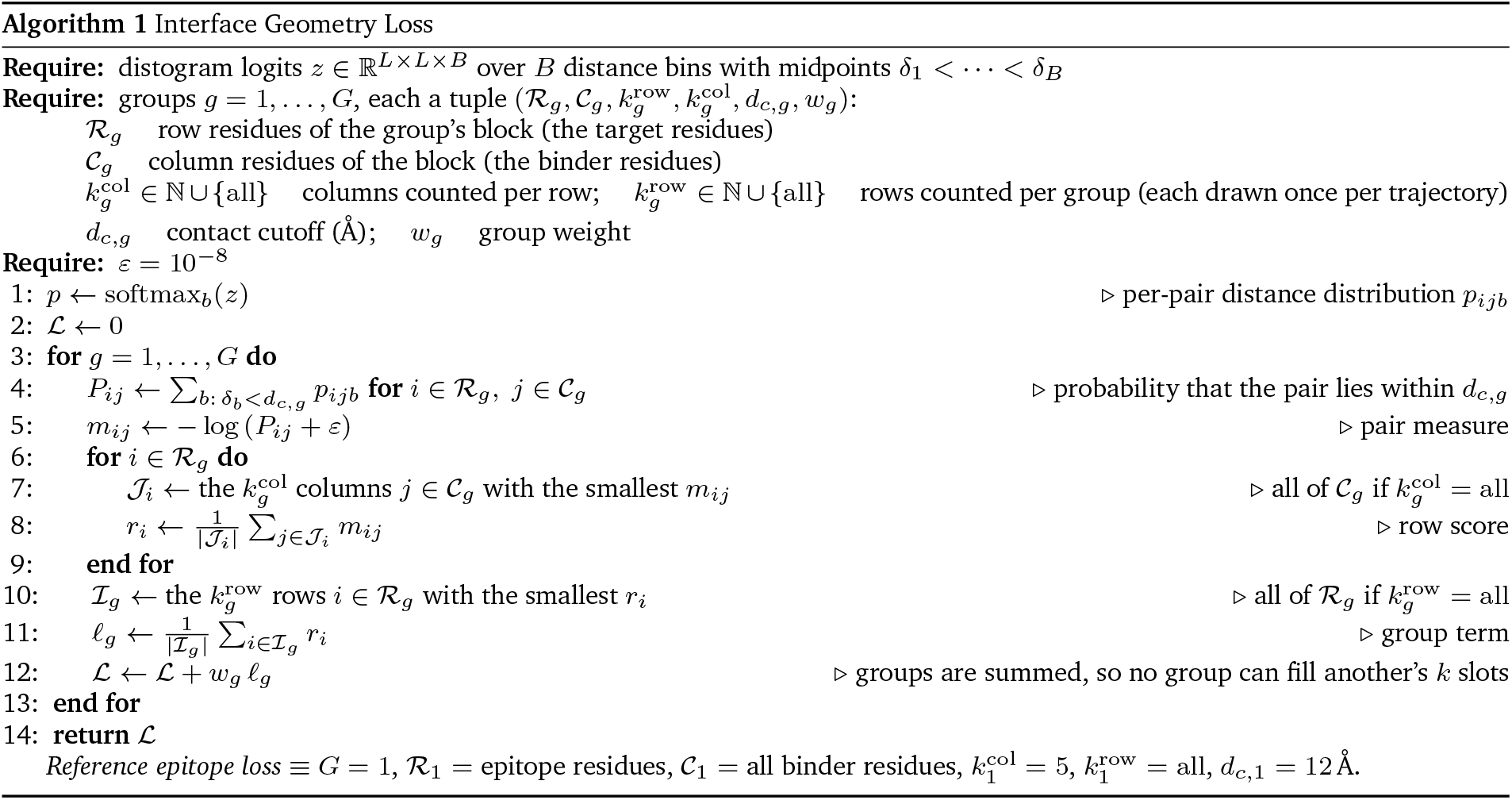
Pseudocode for the proposed interface geometry loss, generalizing the epitope loss of Candido et al. [55] to arbitrarily many interface groups realized as pairs of subsets of rows and columns within the larger distogram, with each containing distinct distance cutoffs and row- and column-count parameters 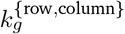.

#### Algorithm 1 Interface Geometry Loss

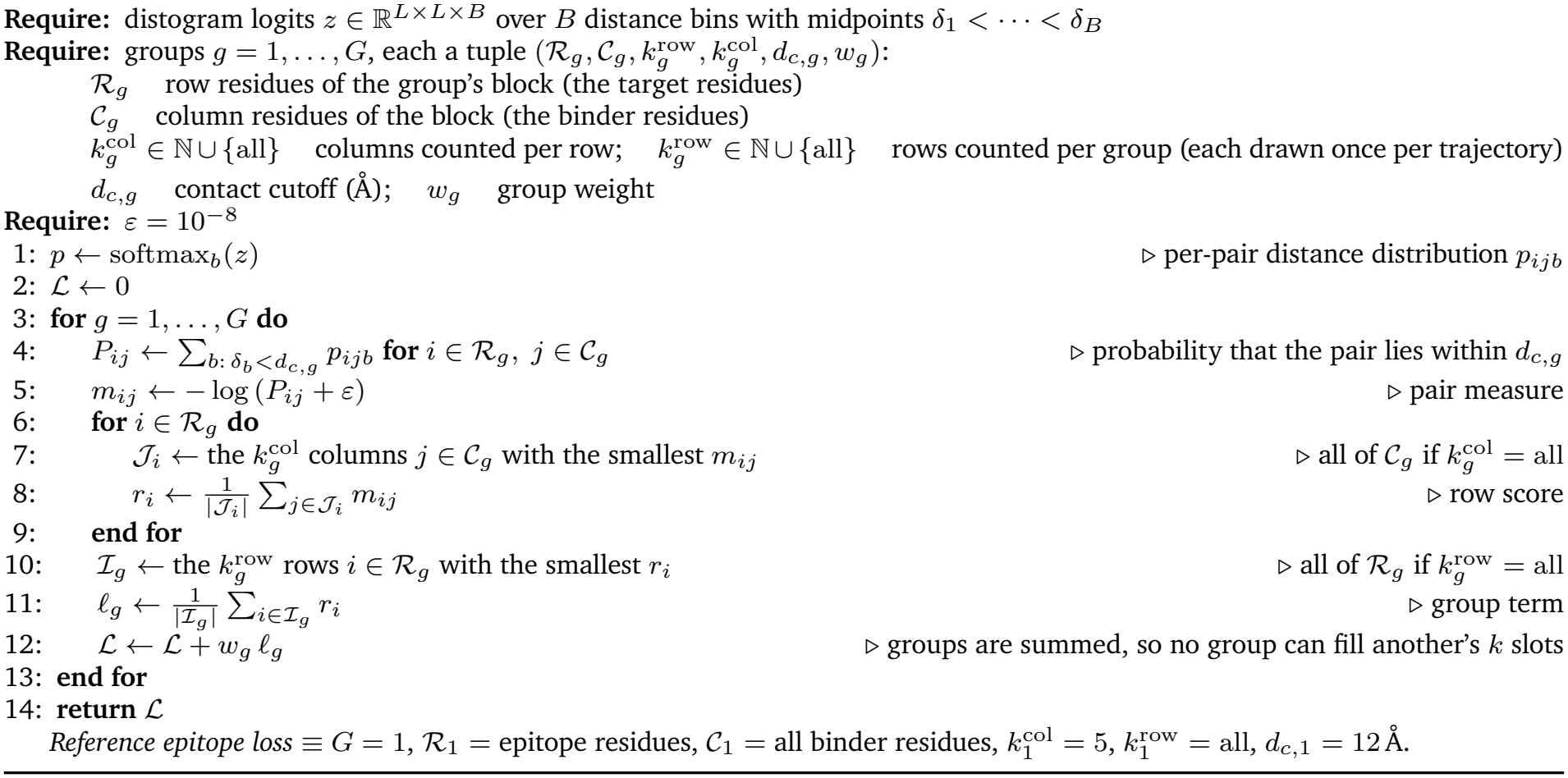

## D Experimental methods

### Protein expression and purification

VHH designs were expressed by cell-free protein synthesis (Gen-Script CFXpress PLUS *E. coli* CFPS kit, cat. RP-D00006-1) from 125 ng of linear DNA template carrying a T7 promoter, ribosome-binding site, the VHH with a C-terminal Twin-Strep tag, a stop codon and a T7 terminator, in the kit’s reaction buffer according to the manufacturer’s instructions. For the screens, reactions were used unpurified after dilution in PBS or Gator K buffer. For purification, variants were captured from the reactions on MagStrep Strep-Tactin XT beads (IBA Lifesciences, cat. 2-5090-010), washed with Buffer W (100 mM Tris-HCl pH 8.0, 150 mM NaCl, 1 mM EDTA) and eluted with Buffer BXT (Buffer W with 50 mM biotin). Protein concentrations were measured on a Qubit Flex fluorometer.

### Bio-layer interferometry

All measurements were made on a Gator Plus instrument (Gator Bio) at 30 ^*°*^C with shaking at 1000 rpm. Unless otherwise noted, all dilutions and buffer steps used Gator K buffer.

### Competitive disruption assay, screen

Soluble His-tagged Fc*ε*RI*α* ectodomain (Sino 13193-H08H-100) was loaded onto anti-His XT probes (Gator Bio) for 90 s, followed by 30 s of buffer. IgE (Abcam AB65866-1008) reconstituted to 100 nM was then bound to the receptor for 120 s, followed by 30 s of buffer. The probe was then moved for 120 s into the cell-free expression reaction of each design (diluted 3*×* in PBS).

### Competitive disruption assay, purified variants

Eight active designs, alongside sdAb026 and Y112A, were purified as above. They were tested at 25, 50, 100, 200 and 400 nM with omalizumab (MedChem-Express HY-P9950) in duplicate. The loading sequence was as in the screen, with phase boundaries at 0, 30, 90, 120, 150 and 180 s, followed by a 600 s exposure to the purified test article.

### Analysis of the disruption traces

Each trace was decomposed into stacked receptor, IgE and VHH components. Let *R*(*t*) be the raw response and *R*_0_ the mean of the last ten reads before receptor loading. The receptor component was fitted on the buffer step after receptor loading, [*t*_1_, *t*_2_], and extrapolated over the rest of the trace:

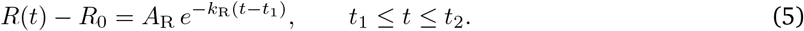

The IgE component is the response above it, 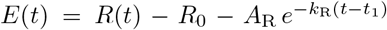, and its decay in the buffer step after IgE loading, [*t*_3_, *t*_4_], was fitted in the same way, 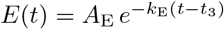, giving *Ê*(*t*), the IgE expected without a disruptor. On exposure to the VHH from *t*_4_, the rise to the first maximum of the trace, at *t*_p_, was assigned to VHH and non-specific binding and carried forward as the constant

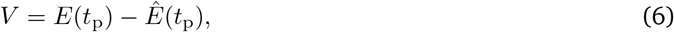

so that any decline below it reports IgE leaving the receptor. The fraction of IgE remaining, the IgE lost at the end of the exposure *t*_end_, and the apparent IgE off-rate *k*_app_ are then

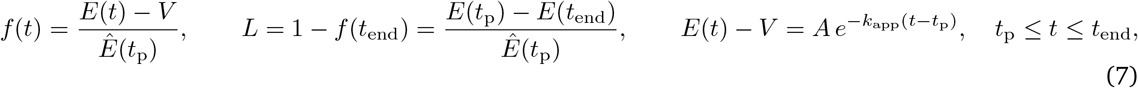

with *f* (*t*_p_) = 1. In the screen, *t*_1_, …, *t*_4_ = 150, 180, 300, 330 s and *t*_end_ = 450 s. A design was called disruptive if

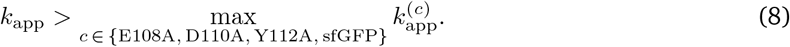

For the purified variants the same decomposition was used, adapted to the missing final buffer step, and the fraction of IgE bound was read as *f* (*t*) at 600 s of exposure.

### Binding assays

VHHs were captured on Strep-Tactin XT probes (Gator Bio) for 60 s from cell-free reaction mix, followed by 60 s of buffer, and exposed to IgE (Abcam AB65866-1008). The disruptor designs were exposed to 100 nM IgE for 120 s with 180 s of dissociation; the anchor designs were exposed to 560 nM for 180 s with 180 s of dissociation. Each trace was referenced to the mean of the last ten reads of the baseline before association, and *R*_max_ is the peak response in the association phase. Each phase was fitted separately with a single exponential, 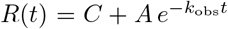 for association and 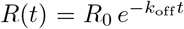 for dissociation over the complete phase, with *t* measured from the start of each phase. For 1:1 binding at a single analyte concentration, *k*_obs_ = *k*_on_[IgE] + *k*_off_, so the apparent association rate constant is

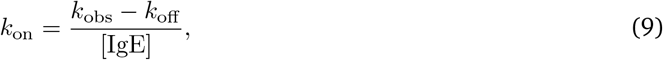

and *K*_D_ = *k*_off_ */k*_on_, reported only where both rates were resolved. Where *k*_off_ falls at the lower bound of the fit it is reported as not resolved.

## E Extended characterization of disruptor designs

Table 4 collects, for every disruptor design and control, the results of the initial disruptor screen and a binding assay against free IgE (Appendix D). From the competitive disruption assay, we report the fraction of receptor-bound IgE lost by the end of the exposure (Equations (7) and (8)). From the binding assay of free IgE at 100 nM, we report the apparent rates *k*_obs_, *k*_off_ and *k*_on_ (Equation (9)).

**Table 4.**
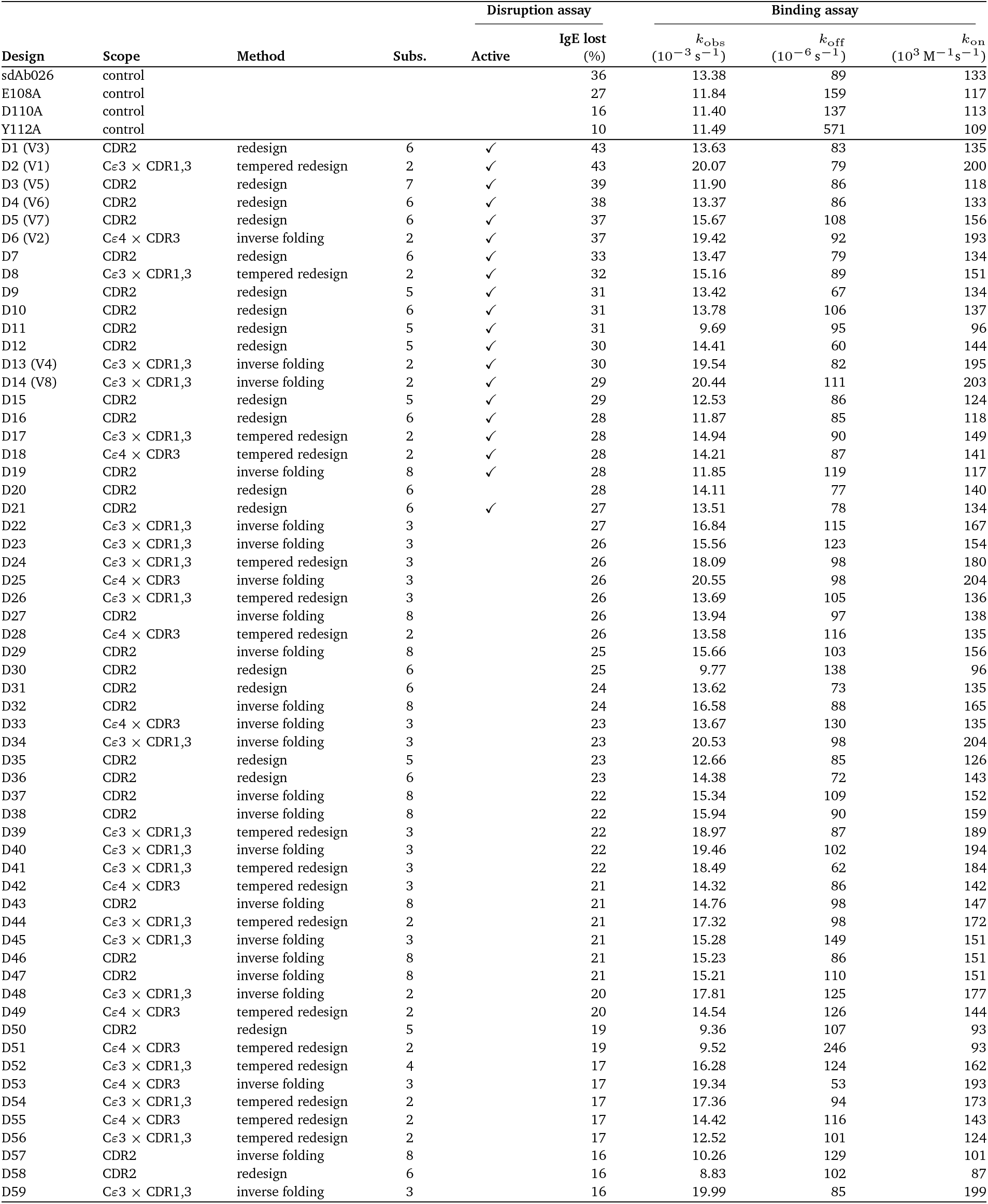
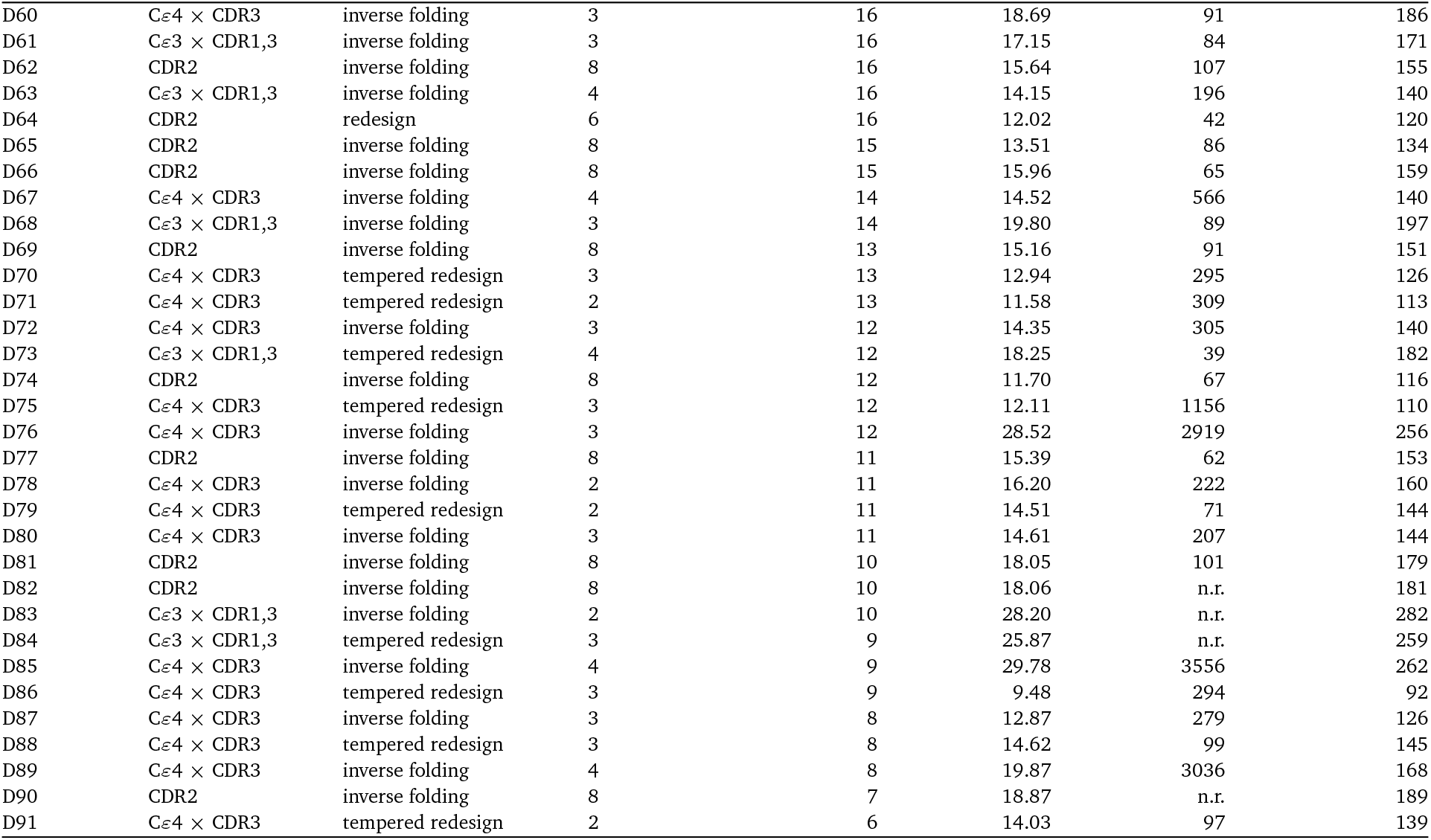
Disruption and binding assays of the disruptor designs and controls. Variants are called active in the disruption assay if they have an apparent IgE off-rate faster than that of every negative control; our primary endpoint (reported in the main text) is the percentage of receptor-bound IgE removed by the end of the exposure. For the binding assay, we report *k*_obs_ and *k*_off_ from single-exponential fits of the association and dissociation phases, and *k*_on_ from Equation (9). n.r., not resolved (rate at the bound of the fit). Scope and method as in Section 3.2; Subs., substitutions relative to sdAb026. Designs D1–D91 are ordered by IgE lost, highest first, and V1–V8 mark the purified variants of Figure 5F–K.

| Design | Scope | Method | Subs. | Disruption assay |  | Binding assay |  |  |
|---|---|---|---|---|---|---|---|---|
| Design | Scope | Method | Subs. | Active | IgE lost<br>(%) | *k*<sub>obs</sub><br>(10<sup>−3</sup> s<sup>−1</sup>) | *k*<sub>off</sub><br>(10<sup>−6</sup> s<sup>−1</sup>) | *k*<sub>on</sub><br>(10<sup>3</sup> M<sup>−1</sup>s<sup>−1</sup>) |
| sdAb026 | control |  |  |  | 36 | 13.38 | 89 | 133 |
| E108A | control |  |  |  | 27 | 11.84 | 159 | 117 |
| D110A | control |  |  |  | 16 | 11.40 | 137 | 113 |
| Y112A | control |  |  |  | 10 | 11.49 | 571 | 109 |
| D1 (V3) | CDR2 | redesign | 6 | ✓ | 43 | 13.63 | 83 | 135 |
| D2 (V1) | Cε3 × CDR1,3 | tempered redesign | 2 | ✓ | 43 | 20.07 | 79 | 200 |
| D3 (V5) | CDR2 | redesign | 7 | ✓ | 39 | 11.90 | 86 | 118 |
| D4 (V6) | CDR2 | redesign | 6 | ✓ | 38 | 13.37 | 86 | 133 |
| D5 (V7) | CDR2 | redesign | 6 | ✓ | 37 | 15.67 | 108 | 156 |
| D6 (V2) | Cε4 × CDR3 | inverse folding | 2 | ✓ | 37 | 19.42 | 92 | 193 |
| D7 | CDR2 | redesign | 6 | ✓ | 33 | 13.47 | 79 | 134 |
| D8 | Cε3 × CDR1,3 | tempered redesign | 2 | ✓ | 32 | 15.16 | 89 | 151 |
| D9 | CDR2 | redesign | 5 | ✓ | 31 | 13.42 | 67 | 134 |
| D10 | CDR2 | redesign | 6 | ✓ | 31 | 13.78 | 106 | 137 |
| D11 | CDR2 | redesign | 5 | ✓ | 31 | 9.69 | 95 | 96 |
| D12 | CDR2 | redesign | 5 | ✓ | 30 | 14.41 | 60 | 144 |
| D13 (V4) | Cε3 × CDR1,3 | inverse folding | 2 | ✓ | 30 | 19.54 | 82 | 195 |
| D14 (V8) | Cε3 × CDR1,3 | inverse folding | 2 | ✓ | 29 | 20.44 | 111 | 203 |
| D15 | CDR2 | redesign | 5 | ✓ | 29 | 12.53 | 86 | 124 |
| D16 | CDR2 | redesign | 6 | ✓ | 28 | 11.87 | 85 | 118 |
| D17 | Cε3 × CDR1,3 | tempered redesign | 2 | ✓ | 28 | 14.94 | 90 | 149 |
| D18 | Cε4 × CDR3 | tempered redesign | 2 | ✓ | 28 | 14.21 | 87 | 141 |
| D19 | CDR2 | inverse folding | 8 | ✓ | 28 | 11.85 | 119 | 117 |
| D20 | CDR2 | redesign | 6 |  | 28 | 14.11 | 77 | 140 |
| D21 | CDR2 | redesign | 6 | ✓ | 27 | 13.51 | 78 | 134 |
| D22 | Cε3 × CDR1,3 | inverse folding | 3 |  | 27 | 16.84 | 115 | 167 |
| D23 | Cε3 × CDR1,3 | inverse folding | 3 |  | 26 | 15.56 | 123 | 154 |
| D24 | Cε3 × CDR1,3 | tempered redesign | 3 |  | 26 | 18.09 | 98 | 180 |
| D25 | Cε4 × CDR3 | inverse folding | 3 |  | 26 | 20.55 | 98 | 204 |
| D26 | Cε3 × CDR1,3 | tempered redesign | 3 |  | 26 | 13.69 | 105 | 136 |
| D27 | CDR2 | inverse folding | 8 |  | 26 | 13.94 | 97 | 138 |
| D28 | Cε4 × CDR3 | tempered redesign | 2 |  | 26 | 13.58 | 116 | 135 |
| D29 | CDR2 | inverse folding | 8 |  | 25 | 15.66 | 103 | 156 |
| D30 | CDR2 | redesign | 6 |  | 25 | 9.77 | 138 | 96 |
| D31 | CDR2 | redesign | 6 |  | 24 | 13.62 | 73 | 135 |
| D32 | CDR2 | inverse folding | 8 |  | 24 | 16.58 | 88 | 165 |
| D33 | Cε4 × CDR3 | inverse folding | 3 |  | 23 | 13.67 | 130 | 135 |
| D34 | Cε3 × CDR1,3 | inverse folding | 3 |  | 23 | 20.53 | 98 | 204 |
| D35 | CDR2 | redesign | 5 |  | 23 | 12.66 | 85 | 126 |
| D36 | CDR2 | redesign | 6 |  | 23 | 14.38 | 72 | 143 |
| D37 | CDR2 | inverse folding | 8 |  | 22 | 15.34 | 109 | 152 |
| D38 | CDR2 | inverse folding | 8 |  | 22 | 15.94 | 90 | 159 |
| D39 | Cε3 × CDR1,3 | tempered redesign | 3 |  | 22 | 18.97 | 87 | 189 |
| D40 | Cε3 × CDR1,3 | inverse folding | 3 |  | 22 | 19.46 | 102 | 194 |
| D41 | Cε3 × CDR1,3 | tempered redesign | 3 |  | 22 | 18.49 | 62 | 184 |
| D42 | Cε4 × CDR3 | tempered redesign | 3 |  | 21 | 14.32 | 86 | 142 |
| D43 | CDR2 | inverse folding | 8 |  | 21 | 14.76 | 98 | 147 |
| D44 | Cε3 × CDR1,3 | tempered redesign | 2 |  | 21 | 17.32 | 98 | 172 |
| D45 | Cε3 × CDR1,3 | inverse folding | 3 |  | 21 | 15.28 | 149 | 151 |
| D46 | CDR2 | inverse folding | 8 |  | 21 | 15.23 | 86 | 151 |
| D47 | CDR2 | inverse folding | 8 |  | 21 | 15.21 | 110 | 151 |
| D48 | Cε3 × CDR1,3 | inverse folding | 2 |  | 20 | 17.81 | 125 | 177 |
| D49 | Cε4 × CDR3 | tempered redesign | 2 |  | 20 | 14.54 | 126 | 144 |
| D50 | CDR2 | redesign | 5 |  | 19 | 9.36 | 107 | 93 |
| D51 | Cε4 × CDR3 | tempered redesign | 2 |  | 19 | 9.52 | 246 | 93 |
| D52 | Cε3 × CDR1,3 | tempered redesign | 4 |  | 17 | 16.28 | 124 | 162 |
| D53 | Cε4 × CDR3 | inverse folding | 3 |  | 17 | 19.34 | 53 | 193 |
| D54 | Cε3 × CDR1,3 | tempered redesign | 2 |  | 17 | 17.36 | 94 | 173 |
| D55 | Cε4 × CDR3 | tempered redesign | 2 |  | 17 | 14.42 | 116 | 143 |
| D56 | Cε3 × CDR1,3 | tempered redesign | 2 |  | 17 | 12.52 | 101 | 124 |
| D57 | CDR2 | inverse folding | 8 |  | 16 | 10.26 | 129 | 101 |
| D58 | CDR2 | redesign | 6 |  | 16 | 8.83 | 102 | 87 |
| D59 | Cε3 × CDR1,3 | inverse folding | 3 |  | 16 | 19.99 | 85 | 199 |
| D60 | Cε4 × CDR3 | inverse folding | 3 |  | 16 | 18.69 | 91 | 186 |
| D61 | Cε3 × CDR1,3 | inverse folding | 3 |  | 16 | 17.15 | 84 | 171 |
| D62 | CDR2 | inverse folding | 8 |  | 16 | 15.64 | 107 | 155 |
| D63 | Cε3 × CDR1,3 | inverse folding | 4 |  | 16 | 14.15 | 196 | 140 |
| D64 | CDR2 | redesign | 6 |  | 16 | 12.02 | 42 | 120 |
| D65 | CDR2 | inverse folding | 8 |  | 15 | 13.51 | 86 | 134 |
| D66 | CDR2 | inverse folding | 8 |  | 15 | 15.96 | 65 | 159 |
| D67 | Cε4 × CDR3 | inverse folding | 4 |  | 14 | 14.52 | 566 | 140 |
| D68 | Cε3 × CDR1,3 | inverse folding | 3 |  | 14 | 19.80 | 89 | 197 |
| D69 | CDR2 | inverse folding | 8 |  | 13 | 15.16 | 91 | 151 |
| D70 | Cε4 × CDR3 | tempered redesign | 3 |  | 13 | 12.94 | 295 | 126 |
| D71 | Cε4 × CDR3 | tempered redesign | 2 |  | 13 | 11.58 | 309 | 113 |
| D72 | Cε4 × CDR3 | inverse folding | 3 |  | 12 | 14.35 | 305 | 140 |
| D73 | Cε3 × CDR1,3 | tempered redesign | 4 |  | 12 | 18.25 | 39 | 182 |
| D74 | CDR2 | inverse folding | 8 |  | 12 | 11.70 | 67 | 116 |
| D75 | Cε4 × CDR3 | tempered redesign | 3 |  | 12 | 12.11 | 1156 | 110 |
| D76 | Cε4 × CDR3 | inverse folding | 3 |  | 12 | 28.52 | 2919 | 256 |
| D77 | CDR2 | inverse folding | 8 |  | 11 | 15.39 | 62 | 153 |
| D78 | Cε4 × CDR3 | inverse folding | 2 |  | 11 | 16.20 | 222 | 160 |
| D79 | Cε4 × CDR3 | tempered redesign | 2 |  | 11 | 14.51 | 71 | 144 |
| D80 | Cε4 × CDR3 | inverse folding | 3 |  | 11 | 14.61 | 207 | 144 |
| D81 | CDR2 | inverse folding | 8 |  | 10 | 18.05 | 101 | 179 |
| D82 | CDR2 | inverse folding | 8 |  | 10 | 18.06 | n.r. | 181 |
| D83 | Cε3 × CDR1,3 | inverse folding | 2 |  | 10 | 28.20 | n.r. | 282 |
| D84 | Cε3 × CDR1,3 | tempered redesign | 3 |  | 9 | 25.87 | n.r. | 259 |
| D85 | Cε4 × CDR3 | inverse folding | 4 |  | 9 | 29.78 | 3556 | 262 |
| D86 | Cε4 × CDR3 | tempered redesign | 3 |  | 9 | 9.48 | 294 | 92 |
| D87 | Cε4 × CDR3 | inverse folding | 3 |  | 8 | 12.87 | 279 | 126 |
| D88 | Cε4 × CDR3 | tempered redesign | 3 |  | 8 | 14.62 | 99 | 145 |
| D89 | Cε4 × CDR3 | inverse folding | 4 |  | 8 | 19.87 | 3036 | 168 |
| D90 | CDR2 | inverse folding | 8 |  | 7 | 18.87 | n.r. | 189 |
| D91 | Cε4 × CDR3 | tempered redesign | 2 |  | 6 | 14.03 | 97 | 139 |

## Footnotes

1 Relative model performance on 7SHT and 5NQW is likely a partial artifact of their deposition dates of October 2021 and January 2018, respectively, and in relation to model training cutoffs.

## References

[1] Scott H. Sicherer and Hugh A. Sampson. Food allergy: A review and update on epidemiology, pathogenesis, diagnosis, prevention, and management. Journal of Allergy and Clinical Immunology, 141 (1):41–58, 2018. doi: 10.1016/j.jaci.2017.11.003.

[2] Ruchi S. Gupta, Elizabeth E. Springston, Manoj R. Warrier, et al. The prevalence, severity, and distribution of childhood food allergy in the United States. Pediatrics, 128(1):e9–e17, 2011. doi: 10.1542/peds.2011-0204.

[3] Ruchi S. Gupta, Christopher M. Warren, Bridget M. Smith, et al. Prevalence and severity of food allergies among US adults. JAMA Network Open, 2(1):e185630, 2019. doi: 10.1001/jamanetworkopen.2018.5630.

[4] Christopher M. Warren, Ozge Nur Aktas, Lorenzo J. Manalo, Tami R. Bartell, and Ruchi S. Gupta. The epidemiology of multifood allergy in the United States: A population-based study. Annals of Allergy, Asthma & Immunology, 130(5):637–648.e5, 2023. doi: 10.1016/j.anai.2022.12.031.

[5] U.S. Food and Drug Administration, Center for Drug Evaluation and Research. NDA/BLA multi-disciplinary review and evaluation: sBLA 103976/S-5245, Xolair (omalizumab). FDA review, BLA 103976, supplement 5245, 2024.

[6] Scott C. Garman, Beth A. Wurzburg, Svetlana S. Tarchevskaya, Jean-Pierre Kinet, and Theodore S. Jardetzky. Structure of the Fc fragment of human IgE bound to its high-affinity receptor FcεRIα. Nature, 406(6793):259–266, 2000. doi: 10.1038/35018500.

[7] Mary D. Holdom, Anna M. Davies, Joanne E. Nettleship, et al. Conformational changes in IgE contribute to its uniquely slow dissociation rate from receptor FcεRI. Nature Structural & Molecular Biology, 18(5):571–576, 2011. doi: 10.1038/nsmb.2044.

[8] Beomkyu Kim, Alexander Eggel, Svetlana S. Tarchevskaya, Monique Vogel, Heino Prinz, and Theodore S. Jardetzky. Accelerated disassembly of IgE-receptor complexes by a disruptive macro-molecular inhibitor. Nature, 491(7425):613–617, 2012. doi: 10.1038/nature11546.

[9] Rodrigo Jiménez-Saiz, Derek K. Chu, Talveer S. Mandur, et al. Lifelong memory responses perpetuate humoral TH2 immunity and anaphylaxis in food allergy. Journal of Allergy and Clinical Immunology, 140(6):1604–1615.e5, 2017. doi: 10.1016/j.jaci.2017.01.018.

[10] Miyo Ota, Kenneth B. Hoehn, Weslley Fernandes-Braga, et al. CD23+IgG1+ memory B cells are poised to switch to pathogenic IgE production in food allergy. Science Translational Medicine, 16 (733):eadi0673, 2024. doi: 10.1126/scitranslmed.adi0673.

[11] Joshua F. E. Koenig, Niels Peter H. Knudsen, Allyssa Phelps, et al. Type 2–polarized memory B cells hold allergen-specific IgE memory. Science Translational Medicine, 16(733):eadi0944, 2024. doi: 10.1126/scitranslmed.adi0944.

[12] Monica G. Lawrence, Judith A. Woodfolk, Alexander J. Schuyler, Leland C. Stillman, Martin D. Chapman, and Thomas A.E. Platts-Mills. Half-life of IgE in serum and skin: Consequences for anti-IgE therapy in patients with allergic disease. Journal of Allergy and Clinical Immunology, 139(2):422–428.e4, 2017. doi: 10.1016/j.jaci.2016.04.056.

[13] Roger M. Cass and Burton R. Andersen. The disappearance rate of skin-sensitizing antibody activity after intradermal administration. Journal of Allergy, 42(1):29–35, 1968. doi: 10.1016/0021-8707(68)90129-9.

[14] Shuichi Kubo, Toshinori Nakayama, Kunie Matsuoka, Hiromichi Yonekawa, and Hajime Karasuyama. Long term maintenance of IgE-mediated memory in mast cells in the absence of detectable serum IgE. The Journal of Immunology, 170(2):775–780, 2003. doi: 10.4049/jimmunol.170.2.775.

[15] Donald W. MacGlashan. Releasability of human basophils: Cellular sensitivity and maximal histamine release are independent variables. Journal of Allergy and Clinical Immunology, 91(2):605–615, 1993. doi: 10.1016/0091-6749(93)90266-i.

[16] Donald W. MacGlashan. Basophil activation testing. Journal of Allergy and Clinical Immunology, 132 (4):777–787, 2013. doi: 10.1016/j.jaci.2013.06.038.

[17] Donald W. MacGlashan and Sarbjit S. Saini. Omalizumab increases the intrinsic sensitivity of human basophils to IgE-mediated stimulation. Journal of Allergy and Clinical Immunology, 132(4): 906–911.e4, 2013. doi: 10.1016/j.jaci.2013.04.056.

[18] Robert A. Wood, Alkis Togias, Scott H. Sicherer, et al. Omalizumab for the treatment of multiple food allergies. New England Journal of Medicine, 390(10):889–899, 2024. doi: 10.1056/NEJMoa2312382.

[19] Genentech, Inc. XOLAIR (omalizumab) injection, for subcutaneous use; XOLAIR (omalizumab) for injection, for subcutaneous use [prescribing information]. U.S. prescribing information, BLA 103976, 2026.

[20] Tse Wen Chang, Frances M. Davis, Nai-Chau Sun, Cecily R. Y. Sun, Donald W. MacGlashan, and Robert G. Hamilton. Monoclonal antibodies specific for human IgE-producing B cells: A potential therapeutic for IgE-mediated allergic diseases. Nature Biotechnology, 8(2):122–126, 1990. doi: 10.1038/nbt0290-122.

[21] L. G. Presta, S. J. Lahr, R. L. Shields, et al. Humanization of an antibody directed against IgE. The Journal of Immunology, 151(5):2623–2632, 1993. doi: 10.4049/jimmunol.151.5.2623.

[22] D. W. MacGlashan, B. S. Bochner, D. C. Adelman, et al. Down-regulation of FcεRI expression on human basophils during in vivo treatment of atopic patients with anti-IgE antibody. The Journal of Immunology, 158(3):1438–1445, 1997. doi: 10.4049/jimmunol.158.3.1438.

[23] John A. Eckman, Patricia M. Sterba, Denise Kelly, et al. Effects of omalizumab on basophil and mast cell responses using an intranasal cat allergen challenge. Journal of Allergy and Clinical Immunology, 125(4):889–895.e7, 2010. doi: 10.1016/j.jaci.2009.09.012.

[24] Lisa A. Beck, Gregory V. Marcotte, Donald MacGlashan, Alkis Togias, and Sarbjit Saini. Omalizumab-induced reductions in mast cell FcεRI expression and function. Journal of Allergy and Clinical Immunology, 114(3):527–530, 2004. doi: 10.1016/j.jaci.2004.06.032.

[25] Luke F. Pennington, Svetlana Tarchevskaya, Daniel Brigger, et al. Structural basis of omalizumab therapy and omalizumab-mediated IgE exchange. Nature Communications, 7(1):11610, 2016. doi: 10.1038/ncomms11610.

[26] Anna M. Davies, Elizabeth G. Allan, Anthony H. Keeble, et al. Allosteric mechanism of action of the therapeutic anti-IgE antibody omalizumab. Journal of Biological Chemistry, 292(24):9975–9987, 2017. doi: 10.1074/jbc.M117.776476.

[27] Alexander Eggel, Günther Baravalle, Gabriel Hobi, et al. Accelerated dissociation of IgE-FcεRI complexes by disruptive inhibitors actively desensitizes allergic effector cells. Journal of Allergy and Clinical Immunology, 133(6):1709–1719.e8, 2014. doi: 10.1016/j.jaci.2014.02.005.

[28] Michael J. Baumann, Alexander Eggel, Patrick Amstutz, Beda M. Stadler, and Monique Vogel. DARPins against a functional IgE epitope. Immunology Letters, 133(2):78–84, 2010. doi: 10.1016/j.imlet.2010.07.005.

[29] A. Eggel, P. Buschor, M. J. Baumann, P. Amstutz, B. M. Stadler, and M. Vogel. Inhibition of ongoing allergic reactions using a novel anti-IgE DARPin-Fc fusion protein. Allergy, 66(7):961–968, 2011. doi: 10.1111/j.1398-9995.2011.02546.x.

[30] Alexander Eggel and Theodore S. Jardetzky. Structural and functional insights into IgE receptor interactions and disruptive inhibition. Immunological Reviews, 331(1):e70031, 2025. doi: 10.1111/imr.70031.

[31] Michael T. Stumpp, Keith M. Dawson, and H. Kaspar Binz. Beyond antibodies: The DARPin® drug platform. BioDrugs, 34(4):423–433, 2020. doi: 10.1007/s40259-020-00429-8.

[32] Luke F. Pennington, Pascal Gasser, Silke Kleinboelting, et al. Directed evolution of and structural insights into antibody-mediated disruption of a stable receptor-ligand complex. Nature Communications, 12(1):7069, 2021. doi: 10.1038/s41467-021-27397-z.

[33] Daniel Brigger, Pascal Guntern, Luke F. Pennington, Robin van Brummelen, Theodore S. Jardetzky, and Alexander Eggel. High-affinity omalizumab variants with optimized disruptive potency prevent anaphylaxis in vivo. Journal of Allergy and Clinical Immunology, 156(5):1290–1302, 2025. doi: 10.1016/j.jaci.2025.05.028.

[34] Excellergy Inc. A phase 1, randomized, double-blind, placebo-controlled, single and multiple ascending dose trial of Exl-111 in healthy participants. ClinicalTrials.gov identifier NCT07356713, 2026.

[35] Peter Meerts, Sigrid Cornelis, Guy Hermans, Jo Vercammen, Frank Kamiel Delphina Verdonck, and Stephanie Staelens. Immunoglobulin single variable domains directed against IgE. Patent WO2012175740A1, December 2012.

[36] M. Rinaldi, T. Denayer, S. Thiolloy, et al. ALX-0962, an anti-IgE Nanobody® with a dual mode of action. European Respiratory Journal 42(Suppl. 57), abstract 1765, 2013.

[37] Peter Meerts, Sigrid Cornelis, Bruno Dombrecht, Jo Vercammen, and Frank Verdonck. Improved polypeptides directed against IgE. Patent WO2014087010A1, June 2014.

[38] Frederic Jabs, Melanie Plum, Nick S. Laursen, et al. Trapping IgE in a closed conformation by mimicking CD23 binding prevents and disrupts FcεRI interaction. Nature Communications, 9(1):7, 2018. doi: 10.1038/s41467-017-02312-7.

[39] Luke F. Pennington, Pascal Gasser, Daniel Brigger, Pascal Guntern, Alexander Eggel, and Theodore S. Jardetzky. Structure-guided design of ultrapotent disruptive IgE inhibitors to rapidly terminate acute allergic reactions. Journal of Allergy and Clinical Immunology, 148(4):1049–1060, 2021. doi: 10.1016/j.jaci.2021.03.050.

[40] Theodore S. Jardetzky, Luke Pennington, Alexander Eggel, and Pascal Gasser. Heteromultimeric proteins for reducing IgE-dependent allergic effector cell activation. Patent WO2022061240A1, March 2022.

[41] Edzard Spillner, Sivelle Coline, Lars Harder Christensen, et al. Displacers of IgE-FcεRI. Patent US12692325B2, July 2026.

[42] Claude Science and Amir Shanehsazzadeh. Autonomous de novo protein binder design with Claude. Anthropic technical report, August 2026. URL https://www-cdn.anthropic.com/30bf50e22a01388bb29bf077ee3f244531594b7a.pdf.

[43] Nick S. Laursen, Frederic Jabs, Edzard Spillner, and Gregers R. Andersen. IgE-Fc in complex with single domain antibody 026. Protein Data Bank entry 5NQW, Worldwide Protein Data Bank, 2017.

[44] Balvinder Dhaliwal, Daopeng Yuan, Marie O. Y. Pang, et al. Crystal structure of IgE bound to its B-cell receptor CD23 reveals a mechanism of reciprocal allosteric inhibition with high affinity receptor FcεRI. Proceedings of the National Academy of Sciences of the United States of America, 109(31):12686–12691, 2012. doi: 10.1073/pnas.1207278109.

[45] Luke F. Pennington and Theodore S. Jardetzky. Structure of a partially disrupted IgE high affinity receptor complex bound to an omalizumab variant. Protein Data Bank entry 7SHT, Worldwide Protein Data Bank, 2021.

[46] Justas Dauparas, Gyu Rie Lee, Robert Pecoraro, et al. Atomic context-conditioned protein sequence design using LigandMPNN. Nature Methods, 22(4):717–723, 2025. doi: 10.1038/s41592-025-02626-1.

[47] Benjamin Fry, Kaia Slaw, and Nicholas F. Polizzi. Zero-shot design of drug-binding proteins via neural iterative selection–expansion. Nature, 656(8126):237–249, 2026. doi: 10.1038/s41586-026-10670-w.

[48] Christopher J. Frank, D. Schiwietz, L. Fuss, Sergey Ovchinnikov, and Hendrik Dietz. AlphaFold2 refinement improves designability of large de novo proteins. bioRxiv, 2024.

[49] Yehlin Cho, Griffin Rangel, Gaurav Bhardwaj, and Sergey Ovchinnikov. Protein Hunter: exploiting structure hallucination within diffusion for protein design. bioRxiv, 2025.

[50] Ali Madani, Ben Krause, Eric R. Greene, et al. Large language models generate functional protein sequences across diverse families. Nature Biotechnology, 41(8):1099–1106, 2023. doi: 10.1038/s41587-022-01618-2.

[51] Thomas Hayes, Roshan Rao, Halil Akin, et al. Simulating 500 million years of evolution with a language model. Science, 387(6736):850–858, 2025. doi: 10.1126/science.ads0018.

[52] Jeffrey A. Ruffolo, Stephen Nayfach, Joseph Gallagher, et al. Design of highly functional genome editors by modelling CRISPR–Cas sequences. Nature, 645(8080):518–525, 2025. doi: 10.1038/s41586-025-09298-z.

[53] Sankar Basu and Björn Wallner. DockQ: a quality measure for protein-protein docking models. PLoS ONE, 11(8):e0161879, 2016. doi: 10.1371/journal.pone.0161879.

[54] Claudio Mirabello and Björn Wallner. DockQ v2: improved automatic quality measure for protein multimers, nucleic acids, and small molecules. Bioinformatics, 40(10):btae586, 2024. doi: 10.1093/bioinformatics/btae586.

[55] Salvatore Candido, Thomas Hayes, Alexander Derry, et al. Language modeling materializes a world model of protein biology. bioRxiv, 2026.

